# Spatiotemporal Autocorrelation of Acoustic Indices: Implications for Biodiversity Monitoring

**DOI:** 10.64898/2026.03.18.712292

**Authors:** Xinyi Jiang, Yixuan Zhang, Zufei Shu, Daiping Wang

**Author notes:** Co-first authors (Xinyi Jiang, Yixuan Zhang).

## Abstract

Passive acoustic monitoring (PAM) is increasingly applied in biodiversity research, yet the spatiotemporal dependence inherent in acoustic indices remains poorly understood, limiting their standardized application and interpretation. We conducted a study using a complete kilometer-grid monitoring system covering an entire protected area (100 grids of 1 × 1 km) in a subtropical region of southern China. We used acoustic recordings from 58 valid monitoring sites to characterize the spatial and temporal autocorrelation of six commonly used acoustic indices. We further examined associations between acoustic indices and environmental characteristics. Spatially, autocorrelation ranges of some acoustic indices extended to approximately 4 km (i.e., the Bioacoustic Index (BIO) and Normalized Difference Soundscape Index (NDSI)). Temporally, all indices showed significant autocorrelation over 2-5 days. Environmental responses were also index-specific, with elevation, distance to roads, and vegetation abundance affecting different indices in contrasting ways. These findings demonstrate that acoustic indices capture spatially and temporally distinct dimensions of soundscape variation rather than a uniform representation of biodiversity. Explicitly accounting for index-specific autocorrelation and environmental responses can improve sampling design and statistical inference in long-term PAM studies.

## 1. Introduction

The accelerating loss of biodiversity requires more efficient monitoring strategies to assess ecological responses to anthropogenic pressures and conservation efforts (Ceballos et al., 2017; Gonzalez et al., 2023). Soundscapes—the aggregate of biological, geophysical, and anthropogenic sounds — offer vital spatiotemporal information on biodiversity dynamics (Ducrettet et al., 2020; Chhaya et al., 2021). In this context, advances in passive acoustic monitoring (PAM) have positioned it as a cornerstone of soundscape-based ecoacoustic research. This automated approach enables the large-scale, long-term collection of cross-taxa ecological data without observer bias or disturbance to wildlife (Ross et al., 2023). PAM has been successfully applied to numerous ecological studies. For example, it has been used to optimize sampling strategies to estimate biodiversity in neotropical forests better (Mauriño et al., 2025); detect seasonal calling patterns and phenology of vocal animals (such as the European green toad (*Bufotes viridis*)) (Kaczmarski et al., 2025); and in marine systems, PAM has revealed rapid ecological responses to harmful algal blooms via changes in fish and marine mammal soundscapes (Rycyk et al., 2020). The flexibility in its sampling design supports multi-scale ecological inference, ranging from individual behavior to community-level metrics (Sugai and Llusia, 2019; Bradfer-Lawrence et al., 2020; Rossetto et al., 2025), establishing PAM as a critical tool for monitoring biodiversity in a rapidly changing world (Sethi et al., 2020).

A primary challenge of PAM, however, is rapidly analyzing vast audio datasets. Variable recording quality, combined with the specialized expertise required for accurate interpretation, complicates the data-processing workflow (Priyadarshani et al., 2018). To address this, acoustic indices offer an alternative for efficiently analyzing large audio datasets. These indices quantify spectral or temporal patterns to characterize the state of acoustic landscapes or communities, reducing the need for manual species identification and intensive computation (Sueur et al., 2014; Farina and James, 2016). Researchers have developed numerous indices using different computational approaches to capture multifaceted acoustic features (Sueur, 2018). Their applications span diverse ecosystems—including forests, freshwater systems, and urban areas—and serve purposes such as wildlife behavior tracking (Rossetto et al., 2025), taxon-specific diversity assessment (Botero-Cañola et al., 2024), habitat quality evaluation (Gómez et al., 2018), climate impact monitoring (Krause and Farina, 2016; McGrann et al., 2022), and detecting human activity encroachment in conservation areas (Sun et al., 2023).

Despite their potential, the broader application of acoustic indices faces persistent challenges in both research validation and practical implementation. In particular, recent studies have questioned their ecological reliability, reporting inconsistent relationships with biodiversity and limited transferability across sites (Giuliani et al., 2024; Sugai et al., 2026). A global meta-analysis revealed only a moderate mean correlation (r = 0.33) between acoustic indices and biodiversity (Alcocer et al., 2022), while empirical studies have reported contrasting relationships between individual indices, such as the Acoustic Complexity Index (ACI), and bird species richness across ecosystems (Myers et al., 2019; McGrann et al., 2022; Dröge et al., 2021). Moreover, recent comparative assessments have highlighted limitations in the design and independent validation of some previous studies, which may contribute to inconsistent evaluations of acoustic-index performance (Bradfer Lawrence et al., 2023). These findings highlight the need to consider not only the ecological meaning of acoustic indices but also the methodological factors that may affect their reliability and transferability across monitoring studies (Botero-Cañola et al., 2024).

A critical yet underexplored methodological factor is the spatiotemporal autocorrelation inherent in acoustic index data. Acoustic monitoring commonly involves multiple recording points distributed across a landscape and continuous or repeated recording through time. Consequently, acoustic index observations from nearby sampling points or from consecutive time intervals are unlikely to be statistically independent. This spatial and temporal dependence is an inherent property of many acoustic datasets, not merely a statistical nuisance to address after data collection. If ignored, spatiotemporal autocorrelation can lead to pseudoreplication, inflated effective sample sizes, underestimated uncertainty, and potentially unreliable statistical inferences (Alston et al., 2023). Therefore, the apparent inconsistency among acoustic index studies may reflect not only differences in ecological relationships or index performance but also how study designs and statistical analyses incorporate the spatial and temporal dependence of acoustic observations.

Despite its potential importance, the intrinsic spatiotemporal dependence of acoustic indices remains poorly characterized. Most studies treat autocorrelation primarily as a confounding factor to be checked before statistical analysis, with spatial autocorrelation often assessed only to determine whether sampling units can be considered independent, rather than to quantify the spatial range over which acoustic similarity persists. Similarly, temporal studies have largely focused on how recording schedules and sampling intervals affect the completeness of ecological information (e.g., Francomano et al., 2021), while comparatively little attention has been given to the temporal persistence of acoustic index time series themselves (Scarpelli et al., 2021; Benocci et al., 2022). Consequently, the spatial and temporal scales over which different acoustic indices remain correlated in natural environments remain insufficiently understood. Systematically characterizing these patterns is therefore essential for developing more robust and efficient acoustic monitoring designs. Quantifying the effective spatial and temporal ranges of acoustic-index autocorrelation could help determine appropriate spacing among recording units, temporal sampling intervals, and the degree of replication needed for statistical independence, reducing pseudoreplication, improving the reliability of ecological inference, and optimizing monitoring resources.

In this study, we conducted passive acoustic monitoring within a coordinated kilometer-grid sampling framework to investigate the spatiotemporal dependence of six commonly used acoustic indices. Specifically, we used continuous long-term acoustic recordings to quantify the spatial autocorrelation and temporal persistence of the six indices and to identify their effective spatial and temporal ranges of dependence. We additionally examined associations between acoustic indices and environment characteristics (vegetation, geography and human disturbance) to provide preliminary ecological context for the observed spatial variation. By integrating spatially structured sampling with continuous temporal monitoring, this study aims to systematically characterize the spatiotemporal autocorrelation of acoustic indices and clarify its implications for sampling design, replication, and statistical inference in PAM-based ecological monitoring. Our findings may provide an empirical basis for developing more efficient and statistically robust monitoring protocols for applying acoustic indices in ecological research.

## 2. Materials and methods

### 2.1. Study site

This work was conducted at the Chebaling National Nature Reserve (114°09’04“-114°16’46“E, 24°40’29“-24°46’21“N), located in Guangdong Province, China. The reserve spans 75.45 square kilometers and lies at the interface between the southern subtropical and middle subtropical climatic zones. Elevation within the reserve range from 330 meters to 1,256 meters, with an average annual temperature of 19.6℃. Annual precipitation varies from 1,150 mm to 2,126 mm (Xu, 1993). The reserve’s primary habitat types are evergreen broad-leaved forests and mixed evergreen coniferous and broad-leaved forests. The reserve is equipped with a comprehensive grid-based monitoring system that covers both the protected area and its surrounding regions. A total of 100 one-kilometer grids (1 km × 1 km) have been established, with 80 located within the reserve (30 in the experimental zone, 28 in the buffer zone, and 22 in the core zone) and 20 situated outside the reserve (Xiao, 2019). For this study, we selected 60 grids from the reserve to ensure representation of key habitat types in the ecological area (Fig. 1). These sampling sites were located in the central and eastern parts of the reserve, with altitudes ranging from 386 meters to 641 meters.

**Fig. 1.**
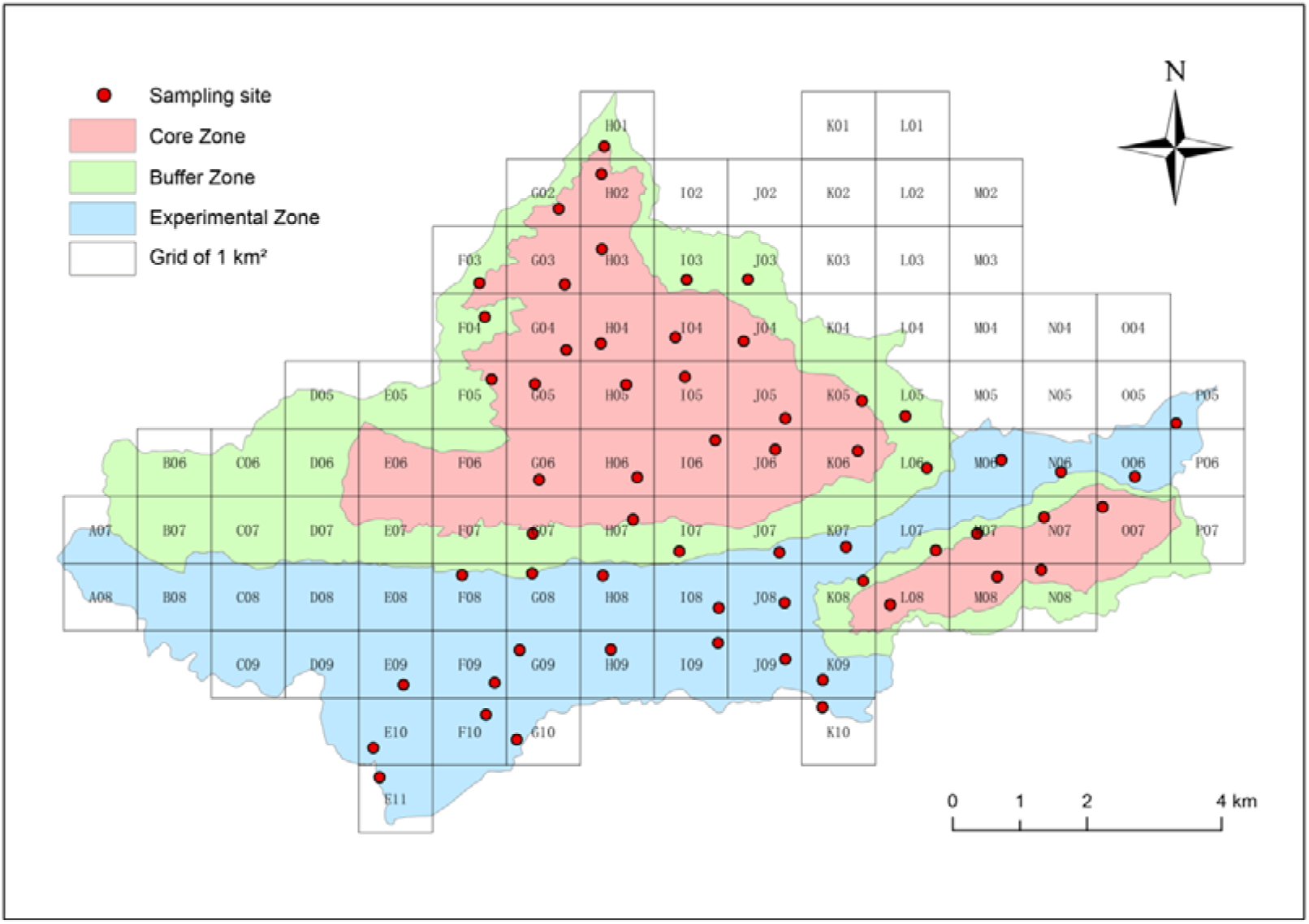
Study area and sampling deployment.

### 2.2 Acoustic monitoring

We established a single sampling site within a 1 km × 1 km grid and equipped it with an automated audio recorder (Song Meter Mini, Wildlife Acoustics). The average distance between two adjacent recording points is 0.975 km. Between August 15, 2022, and January 3, 2023, we conducted three sampling sessions, each lasting at least 38 days. All recording sites remained consistent within each session, ensuring uniformity across sessions within the same batch. To minimize environmental noise interference, we placed the recorders in open areas with sparse vegetation, away from sources of wind and flowing-water noise. We mounted the devices on tree trunks at a height of 3.0-3.5 meters above the ground. The recorders were programmed to operate during the following periods: sunrise (continuous recording from 5:00 to 7:00), sunset (continuous recording from 18:30 to 19:30), and nightly (the first 10 minutes of each hour between 21:00 and 3:00 the following day). Each audio file was limited to 10 minutes, for a total daily recording time of 4 hours. We recorded all audio at 44.1 kHz with 16-bit resolution and saved it in WAV format. In total, we collected 26,124 hours of audio data, amounting to 11.74 TB.

### 2.3 Acoustic indices

We used six commonly applied acoustic indices to characterize the soundscape of the study area. These indices are as follows: 1) The **Acoustic Complexity Index (ACI)** quantifies the variability of sound intensity (amplitude) across adjacent time windows of audio. It estimates the diversity of acoustic communities by measuring fluctuations in sound intensity. A high ACI value indicates increased bird activity, while a low value suggests dominance of continuous insect noise (Pieretti et al., 2011). 2) The **Acoustic Diversity Index (ADI)** measures the complexity of the acoustic composition. It is derived by dividing the frequency spectrum into multiple sub-bands, calculating the proportion of frequency bins exceeding the -50 dBFS threshold within each sub-band, and applying an entropy-based calculation to yield the final ADI value (Villanueva-Rivera et al., 2011). 3) The **Acoustic Evenness Index (AEI)** represents the evenness of sound signal intensity in different frequency bands. The AEI is calculated using the Gini coefficient based on the sound intensity in each 1 kHz frequency band, with values ranging from 0 to 1. High values indicate that acoustic energy is concentrated on a few frequency bands, while low values indicate that a more even distribution of energy across multiple frequency bands or the absence of acoustic activity in all bands (Villanueva-Rivera et al., 2011). 4) The **Bioacoustic Index (BIO)** serves as an important indicator of the abundance of sound-producing species in the environment. It quantifies acoustic complexity by measuring the area of the difference between sound levels at each spectral point and the minimum value (Boelman et al., 2007). 5) The **Acoustic Entropy Index (H)** reflects variations in sound intensity over time and across frequencies. When the acoustic signal is characterized by irregular noise, its value approaches 1, whereas for structured or periodic sounds, it approaches 0. It is calculated by multiplying the time entropy and the spectral entropy (Sueur et al., 2008b). 6) The **Normalized Difference Soundscape Index (NDSI)** quantifies the degree of anthropogenic disturbance by calculating the ratio of anthropophonic to biophonic sounds. Its value ranges from -1 to 1, where values approaching -1 indicate dominance of anthropophonic activity, while values close to 1 reflect the opposite (Kasten et al., 2012).

To minimize the influence of rainfall and equipment noise on the acoustic indices calculations, we first applied a filter to each 10-minute audio file, removing frequency bands below 300 Hz. We then computed the acoustic indices for each recording using the Seewave (Sueur et al., 2008) and Soundecology (Villanueva-Rivera and Pijanowski, 2018) packages in R. The ACI and BIO were calculated over a frequency range of 300 Hz to 12 kHz, while default parameters were used for the computation of all other indices. After calculating the acoustic indices of each audio file, we calculated the daily average of each acoustic index at each site as the basic unit. Because some devices were missing data due to loss or damage, we ultimately retained 58 valid sites, with an average of 112 days of acoustic index recording per site.

### 2.4 Environmental data

We collected environmental data to account for environmental characteristics that may influence acoustic conditions independently of biological acoustic activity. These variables were categorized into three groups: (1) vegetation characteristics, including vegetation species richness and abundance; (2) geographical characteristics, including elevation and slope; and (3) human disturbance, including distance to the nearest road and village.

We obtained vegetation data for each kilometer grid through systematic vegetation surveys conducted within the protected area. Data on vegetation species richness and abundance for the year 2022 were extracted from the online platform of the Chebailing National Nature Reserve (https://cbl.elab.cnic.cn/). This dataset includes survey results from each grid, documenting 365 vegetation species and their respective abundances. We extracted mean elevation and slope for each grid cell from the ASTER GDEM V3 global digital elevation model. Euclidean distances from each acoustic monitoring site to the nearest road and village were calculated using ArcGIS.

### 2.5 Statistical analysis

#### a) Spatial autocorrelation of acoustic indices

We quantified spatial autocorrelation using a multi-layer spatial correlation framework applied consistently to both acoustic indices and biological community composition. For acoustic indices, spatial autocorrelation was quantified by calculating correlations between sites based on their daily mean values across the acoustic monitoring period. We then grouped surrounding grids of each central grid into five concentric spatial layers, with each layer representing a 1-km increase in distance. Within each layer, we selected four neighboring grids (Fig. 2A) and calculated Spearman’s rank correlation coefficients between the focal grid and each neighbor. We used the mean value of these four coefficients to represent the correlation strength of the layer. As a spatial control, we computed the correlation between each focal grid and four randomly selected grids, repeating the random sampling procedure 1,000 times (Fig.2 B).

**Fig. 2.**
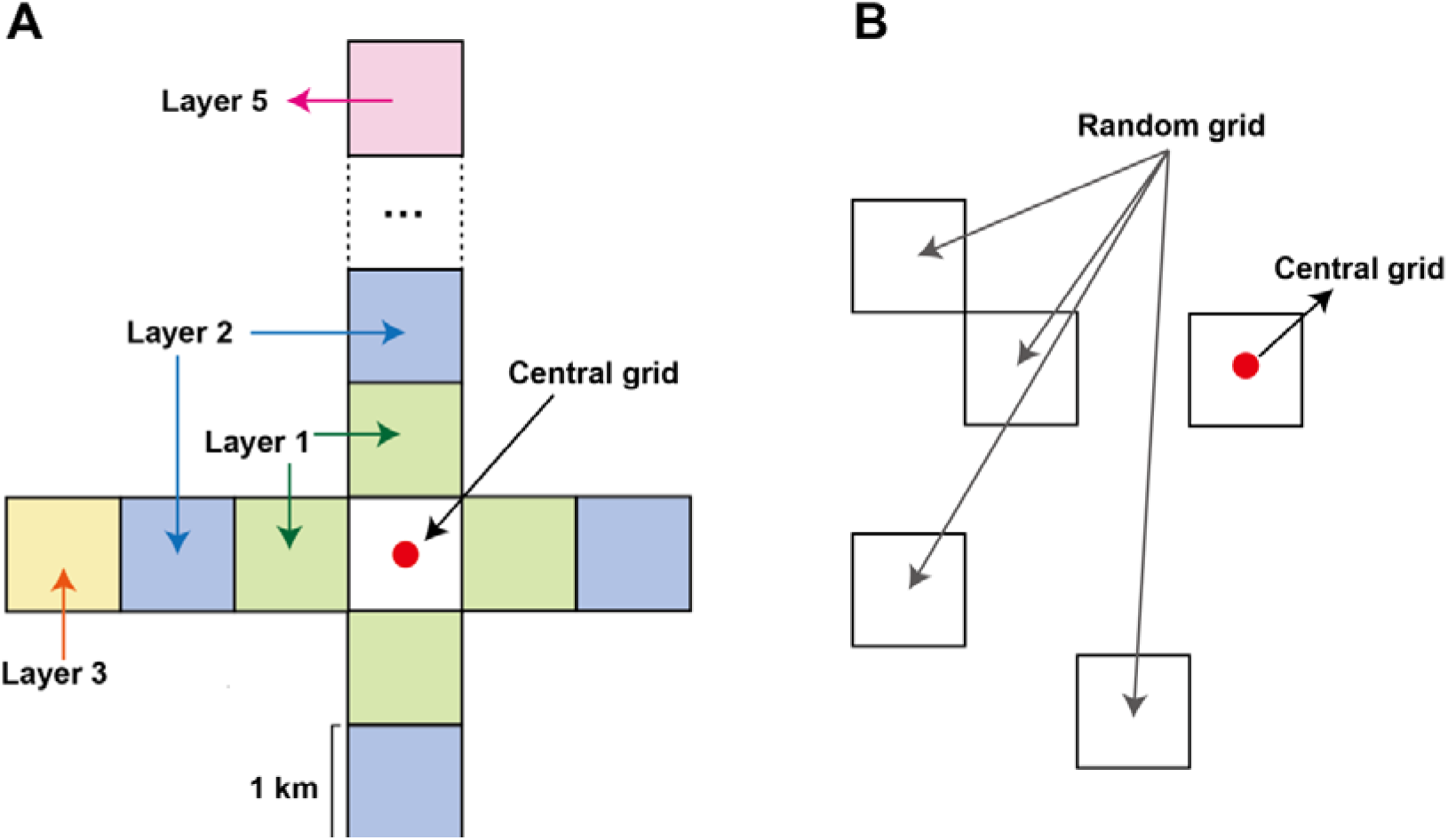
Schematic illustration of grid selection for spatial autocorrelation analysis. (A) Selection of neighboring grids surrounding the central grid. Adjacent grids were grouped into five concentric spatial layers at 1-km distance intervals from the central grid. For each layer, four grids located in the cardinal directions were used to represent spatial relationships at that distance. (B) Construction of the random control group. For each central grid, we randomly selected four grids from the study area to serve as controls. This random sampling procedure was repeated 1,000 times.

We then evaluated differences in correlation coefficients among spatial layers and the random control group using the Kruskal-Wallis rank-sum test, a non-parametric test for detecting differences among multiple groups. When significant differences were detected, we performed Dunn’s post-hoc test to identify which spatial layers differed from the random expectation. To control for multiple comparisons, we adjusted p-values using the Benjamini-Hochberg correction, which controls the false discovery rate. We defined the spatial autocorrelation range as the maximum distance layer whose correlation coefficients remained significantly higher than those of the random control group. For instance, if correlation coefficients for layers 1-3 are consistently and significantly higher than the random expectation, whereas the fourth layer showed no significant difference, we defined the spatial autocorrelation range as extending up to 3 km.

#### b) Temporal autocorrelation of acoustic indices

For temporal autocorrelation analyses, we used the day as the basic temporal unit. We assessed temporal autocorrelation at both the site and the overall levels by using the autocorrelation function (ACF) from the R ‘forecast’ package (Hyndman and Khandakar, 2008). This method measures the correlation between observations in a time series at different time lags, where each lag in this study corresponds to a one-day interval. For each acoustic index, we used the daily mean values recorded over the 112-day sampling period, arranged chronologically by date to match the requirements of time-series analysis. Because the vegetation data lacked repeated temporal measurements, we didn’t analyze vegetation dynamics.

At the site level, we evaluated temporal persistence independently at each monitoring site (n = 58). For each site, we sorted daily mean acoustic indices by sampling time period (112 days). We then calculated the ACF values separately for each site across successive lag orders, where each lag corresponds to one day. Next, we identified the range of significant lags for each time series, which produced one estimate of temporal autocorrelation duration per site.

At the overall level, we summarized the site-level autocorrelation results, resulting in 58 ACF estimates for each lag across the monitoring network. We then calculated the mean ACF value and its 95% confidence interval across sites for each lag order. A lag was considered significant at the overall level if the lower bound of the confidence interval exceeded the threshold 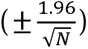, where N represents the sample size, corresponding to the total number of sampling days in this study (N = 112) (Hyndman et al., 2009; Bellisario et al., 2023). For example, if the lower confidence bound remained above the threshold at a lag of 2 days but fell below it at a lag of 3 days, we defined the index’s temporal autocorrelation duration as 2 days.

By combining site-level and overall-level analyses, we assessed both the consistency of temporal persistence among monitoring sites and the regional-scale temporal structure of acoustic indices. This approach allowed us to distinguish persistent temporal patterns that were broadly shared across the monitoring network from those occurring only at individual sites.

#### c) Associations between acoustic indices and environmental characteristics

Before examining the associations between acoustic indices and vegetation diversity, we also evaluated whether vegetation communities exhibited spatial patterns corresponding to those observed in the acoustic indices. Using the same spatial framework described above, we compared vegetation community composition among grids across increasing spatial distances. Because vegetation community composition is multivariate, we used Bray-Curtis (BC) similarity instead of Spearman’s correlation. BC similarity incorporates differences in both species composition and abundance and is widely used to quantify compositional similarity among ecological communities. For each spatial layer, we calculated BC similarity between the focal grid and its four neighboring grids, and used the mean value to represent community similarity at that spatial distance. We then applied the same random-control procedure and statistical comparisons used for acoustic indices. This analysis allowed us to determine whether the spatial scale of vegetation community similarity corresponded to the spatial structure observed in acoustic indices.

We then examined the relationships between acoustic indices and environmental variables using linear models. We modeled each of the six acoustic indices (ACI, ADI, AEI, BIO, H, and NDSI) separately as the response variable. Environmental characteristics were included as explanatory variables, including vegetation species richness, total abundance, elevation, slope, distance to the nearest road, and distance to the nearest village. Before fitting the model, we standardized all response and explanatory variables, allowing us to compare effect sizes among predictors and across acoustic indices. Models were fitted using the lm function in R.

## 3. Results

### a) Spatial Autocorrelation of Acoustic Indices

Correlation analysis and significance tests across hierarchical neighboring grids revealed significant spatial autocorrelation in four acoustic indices. For BIO and NDSI, we detected significant differences from the random control across all four inner layers. For BIO, the adjusted p-values for the first to fourth layers were 0.033, 0.040, 0.040, and 0.041, respectively, whereas for NDSI the adjusted p-values were 0.005 for all four layers. These results indicate high spatial autocorrelation within a radius of about 4 km (Fig.3 A-B). The H showed a significant difference only in the first layer compared with the random control (p-adj = 0.009), indicating that its spatial autocorrelation is confined to approximately 1 km (Fig.3 C). For the ADI, Dunn’s post hoc tests identified a significant difference only between the third spatial layer and the randomly selected control grids (p-adj = 0.041; Fig.3 D). In contrast, correlations observed in the first, second, fourth, and fifth layers did not differ significantly from those of the random controls. This non-monotonic pattern indicates that ADI does not exhibit a clear distance-decay relationship or consistent short-range spatial autocorrelation. Instead, spatial structure in ADI appears weak and scale-specific, with a detectable deviation from randomness occurring only at an intermediate spatial scale of approximately 3 km. Meanwhile, no significant differences from the random control were found in any layer of ACI or AEI, suggesting no clear spatial autocorrelation pattern within the studied extent (Fig.3 E-F, detailed results are provided in the supplementary materials, Table S1).

**Fig. 3.**
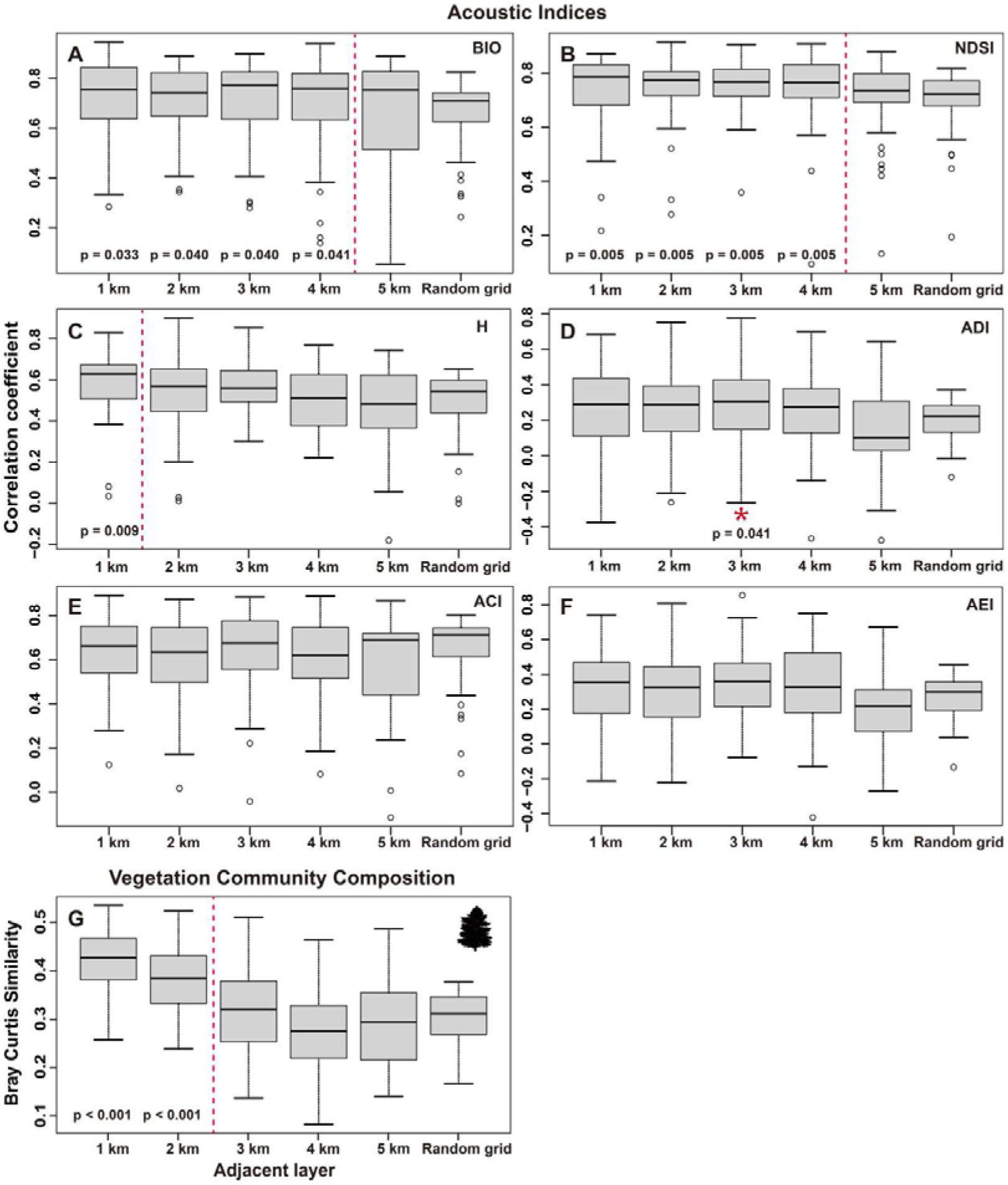
Mean correlations between the acoustic indices of different adjacent layer grids (from the 1st to the 5th order) and the focal central grid, as well as the correlation between random grids and the focal central grid (A-F). Bray-Curtis (BC) similarity of vegetation communities between adjacent layer grids (1st to 5th order) and the focal central grid, compared with BC similarity between random grids and the focal grid (G). The correlation coefficients/similarity of the adjacent layers on the left side of the red dotted line are significantly different from those of the random grids. The correlation coefficient/similarity for this layer differs significantly from that of the random grids, as indicated by the red asterisk. The p-value below the box plot shows the adjusted p-value for the comparison between that layer and the random grids. BIO: Bioacoustic Index; NDSI: Normalized Difference Soundscape Index; H: Acoustic Entropy Index; ADI: Acoustic Diversity Index; ACI: Acoustic Complexity Index; AEI: Acoustic Evenness Index.

Vegetation community composition showed significant spatial autocorrelation within about 2 km (Fig.3 G, detailed results are provided in the supplementary materials, Table S2).

### b) Temporal Autocorrelation of Acoustic Indices

At the site level, the autocorrelation function (ACF) analysis revealed varying degrees of temporal autocorrelation across all six acoustic indices (Fig.4). ACI, ADI, and AEI each exhibited relatively short temporal dependence, each with a median significant lag of 2 days (IQRs: 2-3, 1-3, and 2-4 days, respectively). In contrast, BIO showed a longer median lag of 5 days (IQR: 2-6 days). H and NDSI exhibited intermediate temporal autocorrelation, with median lags of 4 days (IQRs: 3-7 and 2-6 days; detailed results are provided in the supplementary materials, Table S3).

**Fig. 4.**
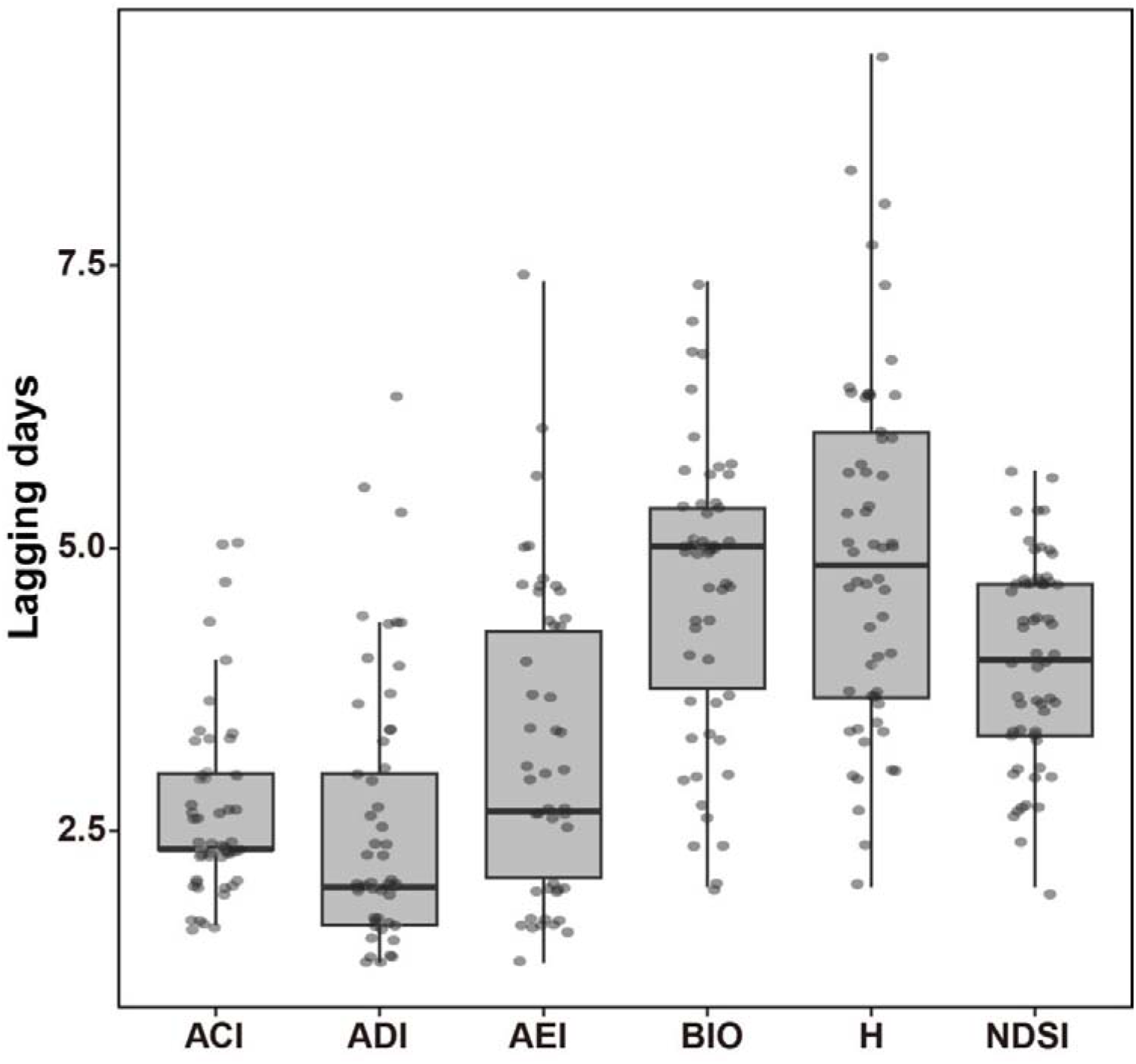
Box plot showing the distribution of temporal autocorrelation duration for daily mean values of each acoustic index at the site level. Points correspond to individual monitoring sites and indicate the lag days at which the autocorrelation of that index’s time series first became non-significant based on autocorrelation function (ACF) analysis. ACI: Acoustic Complexity Index; ADI: Acoustic Diversity Index; AEI: Acoustic Evenness Index; BIO: Bioacoustic Index; H: Acoustic Entropy Index; NDSI: Normalized Difference Soundscape Index.

At the overall level, the significant lag days for ACI, ADI, AEI, and BIO remained consistent with the site-level results (ACI: 2 days; ADI: 2 days; AEI: 2 days; BIO: 5 days; Fig.5 A-D). In contrast, H showed a median lag of 5 days at the overall level, while NDSI showed a reduced median lag of 3 days (Fig.5 E-F, detailed results are provided in the supplementary materials, Table S4).

**Fig. 5.**
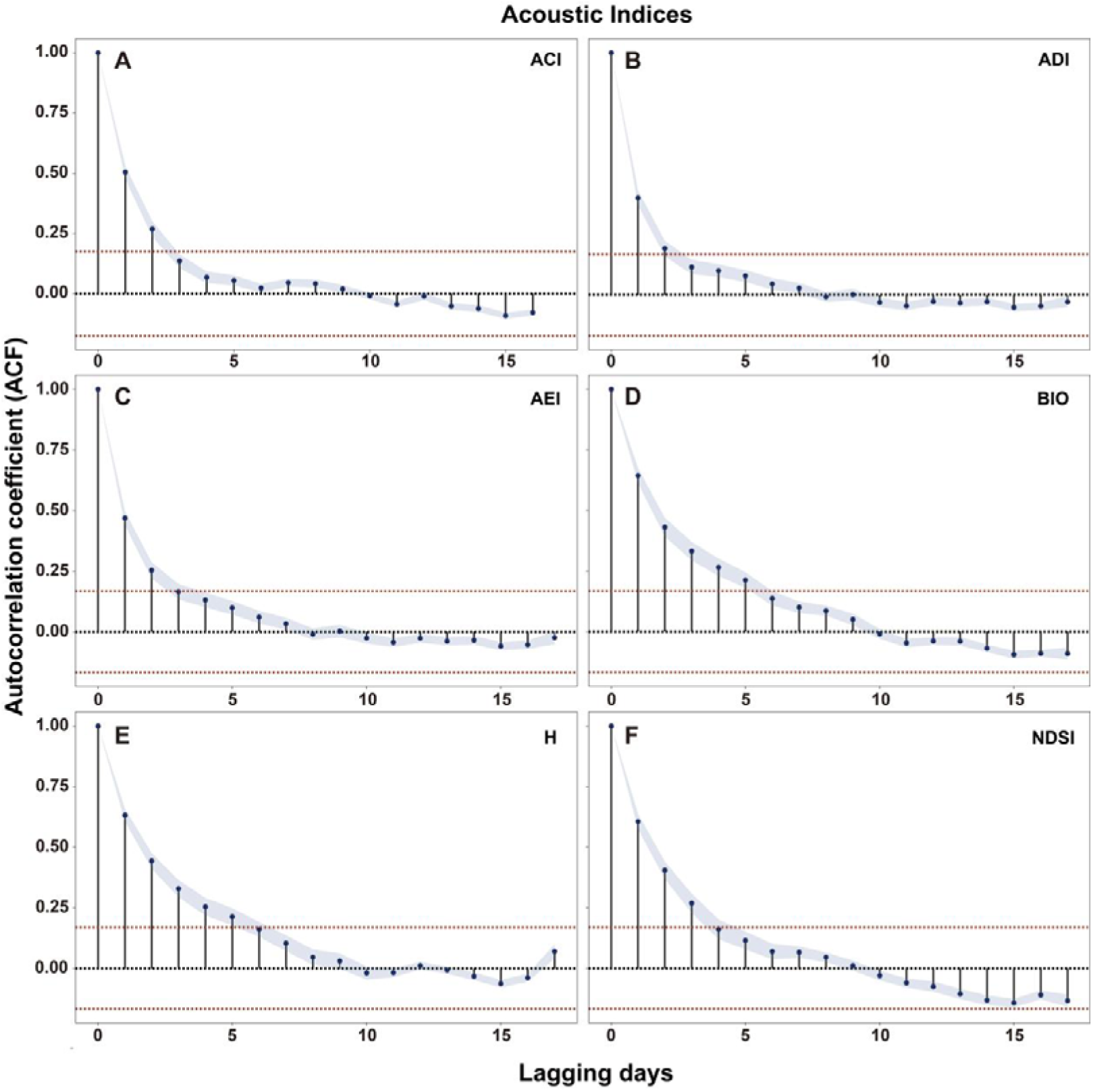
Temporal autocorrelation graphs for each acoustic index (A-F) at the overall level. The horizontal axis denotes the number of lag days, while the vertical axis represents the autocorrelation function value between the corresponding lagged sequence and the original sequence. Values within the bounds of the red dotted lines indicate statistically insignificant correlations. The blue-shaded region corresponds to the confidence interval of the autocorrelation coefficient and only when the lower bound of the 95% confidence interval exceeds the red dotted line is it considered to have a significant autocorrelation. ACI: Acoustic Complexity Index; ADI: Acoustic Diversity Index; AEI: Acoustic Evenness Index; BIO: Bioacoustic Index; H: Acoustic Entropy Index; NDSI: Normalized Difference Soundscape Index.

### c) Associations between acoustic indices and environmental characteristics

Overall, the associations between acoustic indices and environmental variables were generally weak and index-specific, with significant effects detected for vegetation abundance, altitude, and distance to roads in several acoustic indices, whereas most other relationships were not statistically significant.

For ACI, vegetation abundance was positively associated with ACI values (β = 0.398, p = 0.005; Fig.6 A), while no significant effects were detected for other variables. For ADI, none of the environmental variables showed a statistically significant association with the index (Fig.6 B). AEI showed a positive association with elevation (β = 0.540, p = 0.037; Fig.6 C) and a negative association with distance to roads (β = −0.435, p = 0.041). For BIO, elevation was negatively associated with the index (β = −0.784, p = 0.001), whereas distance to roads was positively associated (β = 0.408, p = 0.034; Fig.6 D). H also showed a positive association with distance to roads (β = 0.438, p = 0.028; Fig.6 E), while none of the other environmental variables had significant effects. We detected no significant associations between NDSI and any environmental variables (Fig.6 F, detailed results are provided in the supplementary materials, Table S5).

**Fig. 6.**
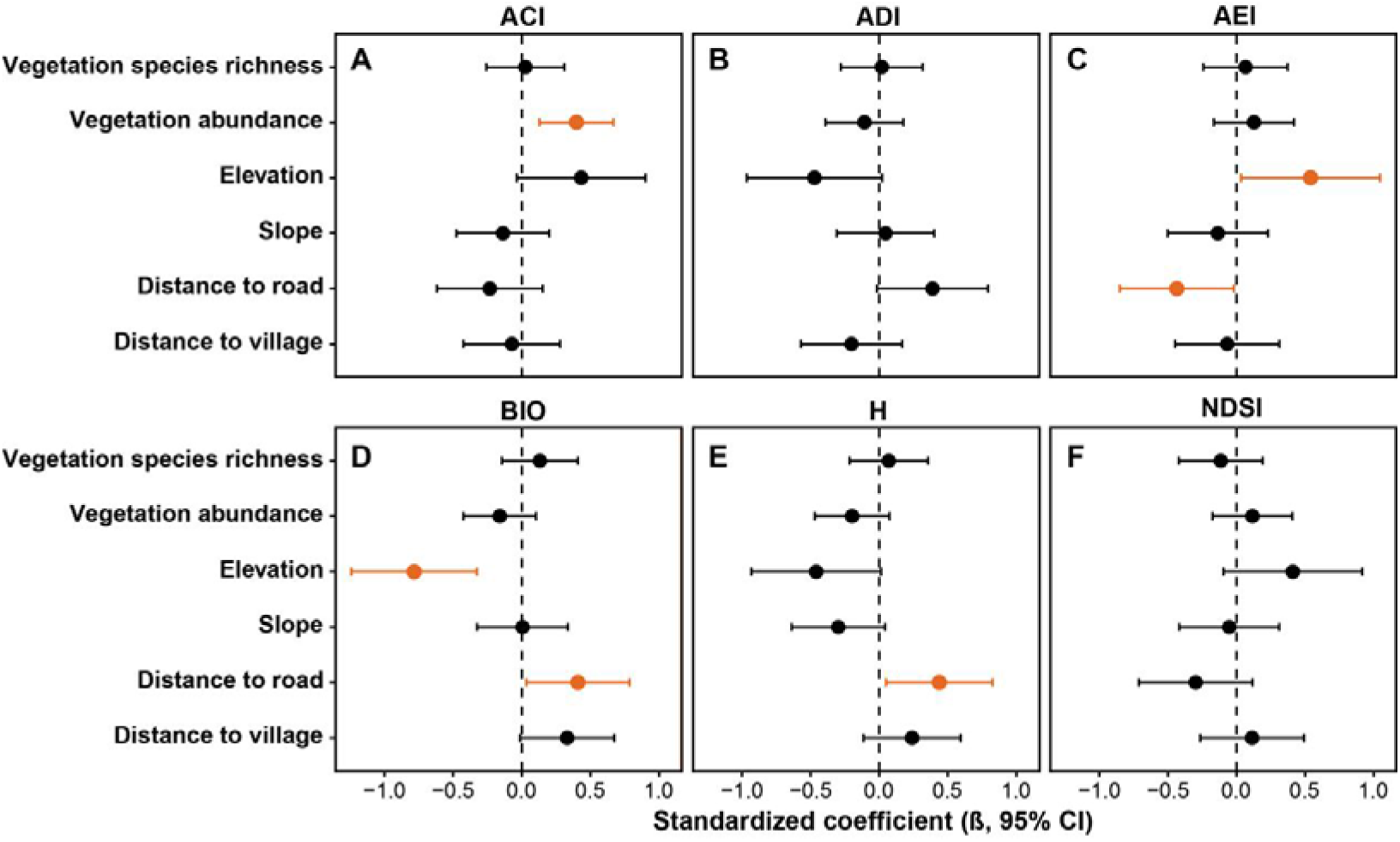
Effect coefficients and 95% confidence intervals of vegetation and environmental variables on acoustic indices based on linear mixed models. Statistically significant effects (95% CI not overlapping zero) are highlighted in orange. ACI: Acoustic Complexity Index; ADI: Acoustic Diversity Index; AEI: Acoustic Evenness Index; BIO: Bioacoustic Index; H: Acoustic Entropy Index; NDSI: Normalized Difference Soundscape Index.

## 4. Discussion

In this study, we systematically quantified the spatial and temporal autocorrelation structures of commonly used acoustic indices based on continuous acoustic monitoring. Our results showed that acoustic indices varied substantially in both spatial correlation ranges and temporal persistence, indicating that different indices represent distinct spatiotemporal dependence regimes. Therefore, the spatiotemporal structure of acoustic indices should be explicitly considered when designing passive acoustic monitoring (PAM) programs and interpreting ecological patterns derived from acoustic data.

### 4.1 Different acoustic indices exhibit distinct spatial autocorrelation scales

We found that spatial autocorrelation ranges differed substantially among acoustic indices, with some indices maintaining significant spatial dependence across several kilometers, while others showed weak or localized spatial structures (Fig.3 A-F). These findings indicate that spatial autocorrelation is not an inherent property of acoustic monitoring systems, but an index-specific characteristic determined by the acoustic properties and ecological processes represented by each metric.

One potential explanation is that different acoustic indices summarize different dimensions of soundscape variation. Indices such as BIO and NDSI showed relatively extensive spatial autocorrelation, with significant spatial dependence extending to approximately 4 km, suggesting that they may capture broad-scale acoustic patterns that are relatively continuous across the landscape. BIO summarizes acoustic energy within a defined frequency range and may therefore integrate the cumulative contribution of multiple sound-producing organisms rather than individual acoustic events. NDSI, in contrast, was designed to quantify the relative contribution of anthropogenic and biological sounds and may respond to spatially extensive disturbance sources such as roads and settlements. More generally, acoustic indices compress complex soundscape information into numerical summaries of different spectral, temporal, or amplitude characteristics, and therefore should not be expected to exhibit identical spatial structures (Bradfer Lawrence et al., 2025)

Differences in the spatial scale and dynamics of sound sources may further contribute to the observed patterns. Biological sounds originate from organisms with different movement capacities, home-range sizes, densities, and calling behaviors, resulting in different spatial footprints within soundscapes. Broadly distributed or temporally coordinated acoustic activity, such as insect choruses or widespread anthropogenic noise, may generate acoustic conditions that remain similar across multiple sampling locations, whereas vocalizations produced by individual animals or small groups may produce more localized spatial variation. Recent work has similarly emphasized that acoustic-index values can reflect mixtures of biotic, anthropogenic, and geophysical sound sources rather than a single ecological component (Mitchell et al., 2020; He et al., 2022; Quinn et al., 2023a). Thus, an index’s spatial autocorrelation scale may partly reflect the dominant spatial scale of the sound sources contributing to it, rather than simply the spatial distribution of biological diversity itself. This may help explain why H showed significant spatial dependence only within approximately 1 km in our study, potentially indicating greater sensitivity to local variation in the distribution or activity of sound-producing organisms than BIO or NDSI.

The spatial patterns of ADI, ACI, and AEI further illustrate that spatial structure cannot be interpreted solely in terms of the geographic distribution of sound sources. ADI showed a weak and non-monotonic spatial pattern, contrasting with the clearer distance-dependent autocorrelation observed for BIO, NDSI, and H. Because ADI describes the evenness of acoustic energy across frequency bands, rather than the overall intensity or temporal variability of acoustic activity, similar ADI values may arise from acoustically different soundscapes, while relatively small changes in particular frequency bands may alter the index independently of overall biological activity. In addition, ADI can be sensitive to background noise and the frequency distribution of acoustic energy, which may weaken a simple distance-dependent spatial pattern (Bradfer Lawrence et al., 2023; Xu et al., 2023). The deviation observed at the third spatial layer may therefore reflect broader-scale habitat heterogeneity or variation in sound-source composition rather than a continuous decline in similarity with geographic distance.

ACI and AEI still showed different spatial patterns. Neither index exhibited significant positive spatial autocorrelation relative to the random controls, suggesting that a simple distance-dependent structure did not characterize their spatial variation. For ACI, the absence of significant spatial dependence across the examined scales is consistent with its sensitivity to temporal fluctuations in acoustic activity. ACI was originally developed to quantify rapid changes in acoustic intensity across successive temporal frames and has subsequently been applied to characterize dynamic biological sound activity (Farina, 2025). Consequently, transient vocalizations or localized acoustic events may generate substantial variation among nearby recording sites and obscure broader spatial continuity. This interpretation does not imply that ACI lacks ecological information; rather, its sensitivity to temporal acoustic variability may make its spatial structure more dependent on the occurrence and distribution of particular sound events than on broad-scale habitat conditions. AEI also showed no significant departure from random expectations across individual distance classes, but the contrast between the inner layers and the outermost layer suggests broader-scale spatial heterogeneity rather than a conventional distance-decay relationship. This pattern indicates that some indices may capture changes between broader landscape contexts rather than gradual changes in acoustic similarity with increasing distance.

Environmental filtering represents another important mechanism shaping the spatial structure of acoustic indices. Vegetation structure, terrain, and habitat characteristics influence sound propagation by affecting attenuation, reflection, and background acoustic conditions (Morton, 1975; Marten and Marler, 1977; Basile et al., 2021). Adjacent habitats often share similar environmental conditions, which can generate spatially structured acoustic patterns even when the biological communities themselves differ. The comparison with vegetation community structure provides an important context for interpreting this relationship. Although vegetation communities exhibited their own spatial autocorrelation in Bray-Curtis similarity, the spatial patterns and scales of acoustic indices did not consistently mirror those of the vegetation community (Fig. 3G). This suggests that the spatial structure of acoustic indices cannot be attributed simply to the spatial distribution of vegetation communities, and that other environmental and acoustic processes may contribute to the observed patterns.

Our environmental models further support this interpretation and reveal that effects were strongly index-specific (Fig.6). Altitude was positively associated with AEI but negatively associated with BIO, whereas distance to roads showed negative associations with AEI but positive associations with BIO and H. ACI was also positively associated with vegetation abundance, while no significant associations with the measured environmental variables were detected for ADI or NDSI. These contrasting responses suggest that environmental gradients do not exert a uniform influence on acoustic indices; instead, their effects depend on which soundscape components each metric emphasizes. For example, topography and vegetation may simultaneously influence habitat conditions, the distribution and activity of sound-producing organisms, and sound transmission, whereas proximity to roads may alter the relative contribution of anthropogenic and biological sounds. Consequently, even when environmental conditions are spatially continuous, their effects may be expressed differently across acoustic metrics, potentially contributing to differences in the extent and strength of spatial autocorrelation observed among indices. Conversely, abrupt changes in habitat structure or landscape configuration may weaken acoustic similarity between nearby sites. Therefore, spatial autocorrelation in acoustic indices should be interpreted as the combined outcome of environmental structure, sound transmission, and the biological and anthropogenic sound sources represented by each index, rather than as a direct representation of spatial patterns in biodiversity.

### 4.2 Temporal persistence differs among acoustic indices

In addition to spatial variation, we found that acoustic indices differed considerably in their temporal autocorrelation duration (Fig.4 and Fig.5). While all indices exhibited significant temporal persistence, the duration of this persistence varied among indices, indicating that different acoustic metrics integrate ecological processes operating at different temporal scales.

Temporal variation in acoustic indices can arise from biological activity patterns. Many organisms exhibit strong daily and seasonal rhythms in vocal activity, such as bird dawn choruses, insect calling activity, and repeated territorial or social vocalizations (Aide et al., 2017; Nascimento et al., 2024). These recurring biological processes can generate short-term temporal continuity in acoustic signals. For indices strongly influenced by transient biological sounds, such as ACI, ADI, and AEI, temporal persistence may reflect repeated daily activity patterns rather than long-lasting ecological states. In contrast, longer temporal autocorrelation observed in indices such as BIO, H, and NDSI may result from more persistent environmental drivers. Abiotic conditions, including humidity, temperature, and wind, influence both biological activity and sound transmission processes, potentially maintaining similar acoustic conditions over several days. Vegetation structure and background acoustic environments often change gradually rather than abruptly, contributing to temporal continuity in acoustic measurements. Such environmental persistence has been recognized as an important factor influencing acoustic index dynamics (Hyland et al., 2023). Anthropogenic factors may also contribute to prolonged temporal dependence. Human activities associated with roads, settlements, and land use often follow predictable schedules and may generate persistent acoustic conditions (Wang et al., 2025). Therefore, indices sensitive to anthropogenic noise, such as NDSI, may retain temporal autocorrelation because of recurring disturbance patterns rather than changes in biological communities.

The observed differences among indices emphasize that temporal autocorrelation should be considered an intrinsic property of individual acoustic metrics. A single universal temporal sampling interval cannot adequately represent the dynamics captured by all acoustic indices.

### 4.3 Implications for PAM sampling design and statistical inference

Our results have important implications for PAM study design and analysis. Because different acoustic indices occupy distinct spatiotemporal dependence regimes, sampling strategies should be determined according to the specific acoustic index and ecological question rather than applying a uniform sampling scale across monitoring programs.

First, spatial sampling design should consider the spatial autocorrelation range of the target acoustic index. When recording units fall within an index’s correlation range, observations may not be independent, potentially inflating effective sample size. Therefore, spatial spacing among recording devices should be selected based on preliminary assessments of index-specific spatial dependence. Second, temporal sampling intervals should account for differences in temporal persistence among indices. For indices with relatively long temporal autocorrelation, frequent sampling may produce highly redundant observations and reduce statistical efficiency. Conversely, indices with short temporal persistence may require higher sampling frequency to capture rapid acoustic changes. Thus, temporal resolution should be matched to the ecological process being investigated. Third, statistical analyses should explicitly incorporate spatiotemporal dependence. Ignoring autocorrelation can underestimate uncertainty, inflate Type I error rates, and bias ecological inference. Appropriate approaches, including spatial random effects, temporal correlation structures, time-series models, or spatiotemporal mixed-effects models, should be considered based on each study’s objectives and data structure.

Overall, our findings suggest that PAM does not operate under a single universal sampling framework. Instead, effective monitoring requires a targeted approach in which the selected acoustic index, research objective, and expected ecological processes jointly determine spatial spacing, temporal resolution, effective sample size, and analytical methods.

### 4.4 Limitations of the study

Several limitations should be considered when interpreting our results. First, acoustic index values are strongly influenced by recording equipment, sampling design, and parameter settings, including frequency ranges and filtering procedures (Bradfer-Lawrence et al., 2019; Hyland et al., 2023; Kemp et al., 2025). Although we followed commonly applied recording protocols and analytical procedures, systematic evaluation of how different methodological choices influence spatiotemporal autocorrelation remains necessary. Second, this study was conducted within a single geographic region and during a limited monitoring period that did not encompass a complete annual phenological cycle. Seasonal changes in biological activity, vegetation dynamics, and environmental conditions may alter the spatiotemporal structure of acoustic indices. Therefore, the autocorrelation ranges identified here should be validated across different ecosystems, climatic regions, and landscape contexts before broad application. Third, although we quantified the spatiotemporal dependence of acoustic indices, we did not directly partition the relative contributions of different sound sources and environmental drivers. Future studies integrating automated species identification, detailed environmental monitoring, and sound-source classification could further clarify the mechanisms generating acoustic autocorrelation patterns.

Despite these limitations, our study provides empirical evidence that acoustic indices have distinct, measurable spatiotemporal structures. Recognizing these index-specific dependence patterns will improve PAM survey design, strengthen statistical inference, and support more reliable ecological interpretation of large-scale acoustic monitoring data.

## 5. Conclusion

Although acoustic indices derived from passive acoustic monitoring (PAM) have increasingly been applied to characterize ecosystem dynamics and biodiversity patterns, their ecological interpretation remains highly context-dependent. In this study, we provide empirical evidence that acoustic indices exhibit distinct and measurable spatiotemporal autocorrelation structures. Our findings demonstrate that acoustic indices should not be treated as a single class of homogeneous ecological metrics, because different indices integrate different acoustic properties, sound sources, and environmental processes operating across different spatial and temporal scales.

The observed variation in autocorrelation patterns highlights that acoustic indices may represent the combined effects of biological activity, environmental filtering, and anthropogenic influences rather than direct measurements of specific ecological components. Therefore, interpreting and applying acoustic indices requires careful consideration of the ecological processes and spatial-temporal scales underlying each metric. Our results further emphasize that no universal sampling framework applies to all PAM studies. Instead, monitoring designs should be tailored to the selected acoustic index and research objectives by considering appropriate spatial spacing, temporal resolution, effective sample size, and statistical approaches for addressing spatiotemporal dependence. By explicitly incorporating index-specific autocorrelation characteristics, future acoustic monitoring studies can improve sampling efficiency, avoid pseudoreplication, and enhance the reliability of ecological inference. Overall, acoustic indices provide valuable complementary information for assessing soundscape variation and ecosystem dynamics. However, their ecological value depends on scale-aware, context-dependent interpretation that integrates acoustic characteristics, environmental conditions, and appropriate analytical frameworks.

## Authors’ contributions

D.W. designed the study; X.J. collected the data and performed the analysis under the supervision of D.W.; X.J., Y.Z., Z.S., and D.W. developed the interpretation of results; X.J. and Y.Z. wrote the first draft under the supervision of D.W., and all authors approved the submission of the current version.

## Acknowledgments

We thank the staff of the Guangdong Chebaling National Nature Reserve Administration for their support in equipment installation and data collection for our research.

## References

Aide, T., Hernández-Serna, A., Campos-Cerqueira, M., Acevedo-Charry, O., Deichmann, J., 2017. Species Richness (of Insects) Drives the Use of Acoustic Space in the Tropics. Remote Sensing 9, 1096. 10.3390/rs9111096

Alcocer, I., Lima, H., Sugai, L.S.M., Llusia, D., 2022. Acoustic indices as proxies for biodiversity: a meta analysis. Biological Reviews 97, 2209–2236. 10.1111/brv.12890

Alston, J.M., Fleming, C.H., Kays, R., Streicher, J.P., Downs, C.T., Ramesh, T., Reineking, B., Calabrese, J.M., 2023. Mitigating pseudoreplication and bias in resource selection functions with autocorrelation informed weighting. Methods Ecol Evol 14, 643–654. 10.1111/2041-210X.14025

Basile, M., Storch, I., Mikusiński, G., 2021. Abundance, species richness and diversity of forest bird assemblages – The relative importance of habitat structures and landscape context. Ecological Indicators 133, 108402. 10.1016/j.ecolind.2021.108402

Bellisario, K., Jessup, L., VanSchaik, J., Dunning, J.B., Graupe, C., Savage, D., Pijanowski, B.C., 2023. Time-series forecasting offers novel quantitative measure to assess loud sound event in an urban park with restored prairie. Ecological Informatics 75, 102100. 10.1016/j.ecoinf.2023.102100

Benocci, R., Roman, H.E., Bisceglie, A., Angelini, F., Brambilla, G., Zambon, G., 2022. Auto-correlations and long time memory of environment sound: The case of an Urban Park in the city of Milan (Italy). Ecological Indicators 134, 108492. 10.1016/j.ecolind.2021.108492

Boelman, N.T., Asner, G.P., Hart, P.J., Martin, R.E., 2007. MULTI TROPHIC INVASION RESISTANCE IN HAWAII: BIOACOUSTICS, FIELD SURVEYS, AND AIRBORNE REMOTE SENSING. Ecological Applications 17, 2137–2144. 10.1890/07-0004.1

Botero-Cañola, S., Wilson, K., Garcia, E., Heisey, M., Reeves, L.E., Burkett-Cadena, N.D., Romagosa, C., Sieving, K.E., Wisely, S.M., 2024. Acoustic indices track local vertebrate biodiversity in a subtropical landscape. Ecological Indicators 166, 112292. 10.1016/j.ecolind.2024.112292

Bradfer-Lawrence, T., Bunnefeld, N., Gardner, N., Willis, S.G., Dent, D.H., 2020. Rapid assessment of avian species richness and abundance using acoustic indices. Ecological Indicators 115, 106400. 10.1016/j.ecolind.2020.106400

Bradfer Lawrence, T., Desjonqueres, C., Eldridge, A., Johnston, A., Metcalf, O., 2023. Using acoustic indices in ecology: Guidance on study design, analyses and interpretation. Methods Ecol Evol 14, 2192–2204. 10.1111/2041-210X.14194

Bradfer Lawrence, T., Duthie, B., Abrahams, C., Adam, M., Barnett, R.J., Beeston, A., Darby, J., Dell, B., Gardner, N., Gasc, A., Heath, B., Howells, N., Janson, M., Kyoseva, M., Luypaert, T., Metcalf, O.C., Nousek McGregor, A.E., Poznansky, F., Ross, S.R.P. J., Sethi, S., Smyth, S., Waddell, E., Froidevaux, J.S.P., 2025. The Acoustic Index User’s Guide: A practical manual for defining, generating and understanding current and future acoustic indices. Methods Ecol Evol 16, 1040–1050. 10.1111/2041-210X.14357

Bradfer Lawrence, T., Gardner, N., Bunnefeld, L., Bunnefeld, N., Willis, S.G., Dent, D.H., 2019. Guidelines for the use of acoustic indices in environmental research. Methods Ecol Evol 10, 1796–1807. 10.1111/2041-210X.13254

Ceballos, G., Ehrlich, P.R., Dirzo, R., 2017. Biological annihilation via the ongoing sixth mass extinction signaled by vertebrate population losses and declines. Proc. Natl. Acad. Sci. U.S.A. 114. 10.1073/pnas.1704949114

Chhaya, V., Lahiri, S., Jagan, M.A., Mohan, R., Pathaw, N.A., Krishnan, A., 2021. Community Bioacoustics: Studying Acoustic Community Structure for Ecological and Conservation Insights. Front. Ecol. Evol. 9, 706445. 10.3389/fevo.2021.706445

Dröge, S., Martin, D.A., Andriafanomezantsoa, R., Burivalova, Z., Fulgence, T.R., Osen, K., Rakotomalala, E., Schwab, D., Wurz, A., Richter, T., Kreft, H., 2021. Listening to a changing landscape: Acoustic indices reflect bird species richness and plot-scale vegetation structure across different land-use types in north-eastern Madagascar. Ecological Indicators 120, 106929. 10.1016/j.ecolind.2020.106929

Ducrettet, M., Forget, P.-M., Ulloa, J.S., Yguel, B., Gaucher, P., Princé, K., Haupert, S., Sueur, J., 2020. Monitoring canopy bird activity in disturbed landscapes with automatic recorders: A case study in the tropics. Biological Conservation 245, 108574. 10.1016/j.biocon.2020.108574

Farina, A., 2025. The acoustic complexity index (ACI): theoretical foundations, applied perspectives and semantics. Oikos 2025, e10760. 10.1111/oik.10760

Farina, A., James, P., 2016. The acoustic communities: Definition, description and ecological role. Biosystems 147, 11–20. 10.1016/j.biosystems.2016.05.011

Francomano, D., Gottesman, B.L., Pijanowski, B.C., 2021. Biogeographical and analytical implications of temporal variability in geographically diverse soundscapes. Ecological Indicators 121, 106794. 10.1016/j.ecolind.2020.106794

Giuliani, M., Mirante, D., Abbondanza, E., Santini, L., 2024. Acoustic indices fail to represent different facets of biodiversity. Ecological Indicators 166, 112451. 10.1016/j.ecolind.2024.112451

Gómez, W.E., Isaza, C.V., Daza, J.M., 2018. Identifying disturbed habitats: A new method from acoustic indices. Ecological Informatics 45, 16–25. 10.1016/j.ecoinf.2018.03.001

Gonzalez, A., Vihervaara, P., Balvanera, P., Bates, A.E., Bayraktarov, E., Bellingham, P.J., Bruder, A., Campbell, J., Catchen, M.D., Cavender-Bares, J., Chase, J., Coops, N., Costello, M.J., Czúcz, B., Delavaud, A., Dornelas, M., Dubois, G., Duffy, E.J., Eggermont, H., Fernandez, M., Fernandez, N., Ferrier, S., Geller, G.N., Gill, M., Gravel, D., Guerra, C.A., Guralnick, R., Harfoot, M., Hirsch, T., Hoban, S., Hughes, A.C., Hugo, W., Hunter, M.E., Isbell, F., Jetz, W., Juergens, N., Kissling, W.D., Krug, C.B., Kullberg, P., Le Bras, Y., Leung, B., Londoño-Murcia, M.C., Lord, J.-M., Loreau, M., Luers, A., Ma, K., MacDonald, A.J., Maes, J., McGeoch, M., Mihoub, J.B., Millette, K.L., Molnar, Z., Montes, E., Mori, A.S., Muller-Karger, F.E., Muraoka, H., Nakaoka, M., Navarro, L., Newbold, T., Niamir, A., Obura, D., O’Connor, M., Paganini, M., Pelletier, D., Pereira, H., Poisot, T., Pollock, L.J., Purvis, A., Radulovici, A., Rocchini, D., Roeoesli, C., Schaepman, M., Schaepman-Strub, G., Schmeller, D.S., Schmiedel, U., Schneider, F.D., Shakya, M.M., Skidmore, A., Skowno, A.L., Takeuchi, Y., Tuanmu, M.-N., Turak, E., Turner, W., Urban, M.C., Urbina-Cardona, N., Valbuena, R., Van De Putte, A., Van Havre, B., Wingate, V.R., Wright, E., Torrelio, C.Z., 2023. A global biodiversity observing system to unite monitoring and guide action. Nat Ecol Evol 7, 1947–1952. 10.1038/s41559-023-02171-0

He, X., Deng, Y., Dong, A., Lin, L., 2022. The relationship between acoustic indices, vegetation, and topographic characteristics is spatially dependent in a tropical forest in southwestern China. Ecological Indicators 142, 109229. 10.1016/j.ecolind.2022.109229

Hyland, E.B., Schulz, A., Quinn, J.E., 2023. Quantifying the Soundscape: How filters change acoustic indices. Ecological Indicators 148, 110061. 10.1016/j.ecolind.2023.110061

Hyndman, R., Athanasopoulos, G., Bergmeir, C., Caceres, G., Chhay, L., Kuroptev, K., O’Hara-Wild, M., Petropoulos, F., Razbash, S., Wang, E., Yasmeen, F., 2009. forecast: Forecasting Functions for Time Series and Linear Models. 10.32614/CRAN.package.forecast

Hyndman, R.J., Khandakar, Y., 2008. Automatic Time Series Forecasting: The **forecast** Package for *R*. J. Stat. Soft. 27. 10.18637/jss.v027.i03

Kaczmarski, M., Kaczmarek, J.M., Radzińska, A., Budka, M., 2025. Passive acoustic monitoring reveals seasonal patterns in European green toad calling activity but fails to accurately reflect population abundance. Sci Rep 15, 26447. 10.1038/s41598-025-11706-3

Kasten, E.P., Gage, S.H., Fox, J., Joo, W., 2012. The remote environmental assessment laboratory’s acoustic library: An archive for studying soundscape ecology. Ecological Informatics 12, 50–67. 10.1016/j.ecoinf.2012.08.001

Krause, B., Farina, A., 2016. Using ecoacoustic methods to survey the impacts of climate change on biodiversity. Biological Conservation 195, 245–254. 10.1016/j.biocon.2016.01.013

Marten, K., Marler, P., 1977. Sound Transmission for Animal Vocalization.

Mauriño, R.A., Zurano, J.P., Zurita, G.A., Di Giacomo, A.S., De Araújo, C.B., 2025. Optimizing passive acoustic monitoring (PAM) for biodiversity studies: Using species–area relationship (SAR) to predict species richness. Ornithological Applications duaf063. 10.1093/ornithapp/duaf063

McGrann, M.C., Wagner, B., Klauer, M., Kaphan, K., Meyer, E., Furnas, B.J., 2022. Using an acoustic complexity index to help monitor climate change effects on avian diversity. Ecological Indicators 142, 109271. 10.1016/j.ecolind.2022.109271

Mitchell, S.L., Bicknell, J.E., Edwards, D.P., Deere, N.J., Bernard, H., Davies, Z.G., Struebig, M.J., 2020. Spatial replication and habitat context matters for assessments of tropical biodiversity using acoustic indices. Ecological Indicators 119, 106717. 10.1016/j.ecolind.2020.106717

Morton, E.S., 1975. Ecological sources of selection on avian sounds. The American Naturalist 109, 17–34.

Myers, D., Berg, H., Maneas, G., 2019. Comparing the soundscapes of organic and conventional olive groves: A potential method for bird diversity monitoring. Ecological Indicators 103, 642–649. 10.1016/j.ecolind.2019.04.030

Nascimento, L.A.D., Pérez-Granados, C., Alencar, J.B.R., Beard, K.H., 2024. Time and habitat structure shape insect acoustic activity in the Amazon.

Pieretti, N., Farina, A., Morri, D., 2011. A new methodology to infer the singing activity of an avian community: The Acoustic Complexity Index (ACI). Ecological Indicators 11, 868–873. 10.1016/j.ecolind.2010.11.005

Priyadarshani, N., Marsland, S., Castro, I., 2018. Automated birdsong recognition in complex acoustic environments: a review. Journal of Avian Biology 49, jav-01447. 10.1111/jav.01447

Quinn, C.A., Burns, P., Hakkenberg, C.R., Salas, L., Pasch, B., Goetz, S.J., Clark, M.L., 2023. Soundscape components inform acoustic index patterns and refine estimates of bird species richness. Front. Remote Sens. 4, 1156837. 10.3389/frsen.2023.1156837

Ross, S.R.P. J., O’Connell, D.P., Deichmann, J.L., Desjonquères, C., Gasc, A., Phillips, J.N., Sethi, S.S., Wood, C.M., Burivalova, Z., 2023. Passive acoustic monitoring provides a fresh perspective on fundamental ecological questions. Functional Ecology 37, 959–975. 10.1111/1365-2435.14275

Rossetto, F., Mathevon, N., Laiolo, P., 2025. Using acoustic indices to detect interspecific bird interactions and behaviour. Ibis ibi.70002. 10.1111/ibi.70002

Rycyk, A.M., Tyson Moore, R.B., Wells, R.S., McHugh, K.A., Berens McCabe, E.J., Mann, D.A., 2020. Passive acoustic listening stations (PALS) show rapid onset of ecological effects of harmful algal blooms in real time. Sci Rep 10, 17863. 10.1038/s41598-020-74647-z

Scarpelli, M.D.A., Liquet, B., Tucker, D., Fuller, S., Roe, P., 2021. Multi-Index Ecoacoustics Analysis for Terrestrial Soundscapes: A New Semi-Automated Approach Using Time-Series Motif Discovery and Random Forest Classification. Front. Ecol. Evol. 9, 738537. 10.3389/fevo.2021.738537

Sethi, S.S., Jones, N.S., Fulcher, B.D., Picinali, L., Clink, D.J., Klinck, H., Orme, C.D.L., Wrege, P.H., Ewers, R.M., 2020. Characterizing soundscapes across diverse ecosystems using a universal acoustic feature set. Proc. Natl. Acad. Sci. U.S.A. 117, 17049–17055. 10.1073/pnas.2004702117

Sueur, J., 2018. Sound Analysis and Synthesis with R. Springer, Switzerland.

Sueur, Jerome, Aubin, T., Simonis, C., 2008. SEEWAVE, A FREE MODULAR TOOL FOR SOUND ANALYSIS AND SYNTHESIS. Bioacoustics 18, 213–226. 10.1080/09524622.2008.9753600

Sueur, J., Farina, A., Gasc, A., Pieretti, N., Pavoine, S., 2014. Acoustic Indices for Biodiversity Assessment and Landscape Investigation. Acta Acustica united with Acustica 100, 772–781. 10.3813/AAA.918757

Sueur, Jérôme, Pavoine, S., Hamerlynck, O., Duvail, S., 2008. Rapid Acoustic Survey for Biodiversity Appraisal. PLoS ONE 3, e4065. 10.1371/journal.pone.0004065

Sugai, L.S.M., Balantic, C., Clink, D.J., Ramesh, V., Kahl, S., Klinck, H., Wood, C.M., 2026. Acoustic indices are not useful for biodiversity research. Methods Ecol Evol 17, 1506–1518. 10.1111/2041-210x.70285

Sugai, L.S.M., Llusia, D., 2019. Bioacoustic time capsules: Using acoustic monitoring to document biodiversity. Ecological Indicators 99, 149–152. 10.1016/j.ecolind.2018.12.021

Sun, Y., Wang, S., Feng, J., Ge, J., Wang, T., 2023. Free-ranging livestock changes the acoustic properties of summer soundscapes in a Northeast Asian temperate forest. Biological Conservation 283, 110123. 10.1016/j.biocon.2023.110123

Villanueva-Rivera, L.J., Pijanowski, B.C., 2018. soundecology: Soundscape Ecology.

Villanueva-Rivera, L.J., Pijanowski, B.C., Doucette, J., Pekin, B., 2011. A primer of acoustic analysis for landscape ecologists. Landscape Ecol 26, 1233–1246. 10.1007/s10980-011-9636-9

Wang, S., Duan, Y., Cao, R., Feng, J., Ge, J., Wang, T., 2025. Road disturbance drives a more simplified soundscape in temperate forests revealed by deep learning and acoustics indices. Biological Conservation 306, 111115. 10.1016/j.biocon.2025.111115

Xiao, Z., 2019. Inventory and Assessment of Wildlife and Its Habitat in Protected Areas—An Example from Chebaling National Nature Reserve, Guangdong, China. China Forestry Publishing House, Beijing.

Xu, Q., 1993. A comprehensive report on investigation in Chebaling National Nature Reserve, in: Collected Papers for Investigation in Chebaling National Nature Reserve. Guangdong Science and Technology Press, Guangzhou, pp. 1–7.

Xu, Z., Chen, L., Pijanowski, B.C., Zhao, Z., 2023. A frequency-dependent acoustic diversity index: A revision to a classic acoustic index for soundscape ecological research. Ecological Indicators 155, 110940. 10.1016/j.ecolind.2023.110940

