## Supplementary material for "Spatiotemporal Autocorrelation of Acoustic Indices: Implications for Biodiversity Monitoring": Table S

* Co-first authors

**Abstract**

Passive acoustic monitoring (PAM) is increasingly applied in biodiversity research, yet the spatial and temporal dependence inherent in acoustic indices remains insufficiently understood, limiting their standardized application and interpretation. We conducted a study using a complete kilometer-grid monitoring system covering an entire protected area (100 grids of 1 × 1 km) in a subtropical region of southern China. We used acoustic recordings from 58 valid monitoring sites to characterize the spatial and temporal autocorrelation of six commonly used acoustic indices. We further examined associations between acoustic indices and environmental characteristics. Spatially, autocorrelation ranges of some acoustic indices extended to approximately 4 km (i.e., the Bioacoustic Index (BIO) and Normalized Difference Soundscape Index (NDSI)). Temporally, all indices showed significant autocorrelation over 2-5 days. Environmental responses were also index-specific, with elevation, distance to roads, and vegetation abundance affecting different indices in contrasting ways. These findings demonstrate that acoustic indices capture spatially and temporally distinct dimensions of soundscape variation rather than a uniform representation of biodiversity. Explicitly accounting for index-specific autocorrelation and environmental responses can improve sampling design and statistical inference in long-term PAM studies.

**Keywords**: soundscape; acoustic indices; spatiotemporal autocorrelation; passive acoustic monitoring

**Table S1.** Results of Dunn’s post-hoc tests comparing spatial autocorrelation across concentric layers against random controls for each acoustic index. Spearman correlation coefficients between the focal grid and each surrounding layer were compared with those from randomly selected grids. p-values were adjusted using the Benjamini-Hochberg method.

| **Acoustic indices** | **Tested layer pairs** | **N** | **Statistic** | **p** | **Adjusted p-value** |
| --- | --- | --- | --- | --- | --- |
| ACI | 1-2 | 55 | -0.699 | 0.485 | 0.791 |
|  | 1-3 | 54 | 0.372 | 0.710 | 0.858 |
|  | 1-4 | 54 | -1.060 | 0.289 | 0.619 |
|  | 1-5 | 47 | -0.974 | 0.330 | 0.619 |
|  | 1-Random grid | 57 | 0.632 | 0.527 | 0.791 |
|  | 2-3 | 54 | 1.058 | 0.290 | 0.619 |
|  | 2-4 | 54 | -0.361 | 0.718 | 0.858 |
|  | 2-5 | 47 | -0.301 | 0.763 | 0.858 |
|  | 2-Random grid | 55 | 1.325 | 0.185 | 0.565 |
|  | 3-4 | 54 | -1.412 | 0.158 | 0.565 |
|  | 3-5 | 47 | -1.316 | 0.188 | 0.565 |
|  | 3-Random grid | 54 | 0.252 | 0.801 | 0.858 |
|  | 4-5 | 47 | 0.047 | 0.963 | 0.963 |
|  | 4-Random grid | 54 | 1.683 | 0.092 | 0.565 |
|  | 5-Random grid | 47 | 1.575 | 0.115 | 0.565 |
| ADI | 1-2 | 55 | -0.418 | 0.676 | 0.887 |
|  | 1-3 | 54 | 0.217 | 0.828 | 0.887 |
|  | 1-4 | 54 | -0.054 | 0.957 | 0.957 |
|  | 1-5 | 47 | -3.210 | 0.001 | 0.009 |
|  | 1-Random grid | 57 | -2.282 | 0.023 | 0.056 |
|  | 2-3 | 54 | 0.628 | 0.530 | 0.795 |
|  | 2-4 | 54 | 0.359 | 0.720 | 0.887 |
|  | 2-5 | 47 | -2.786 | 0.005 | 0.020 |
|  | 2-Random grid | 55 | -1.843 | 0.065 | 0.122 |
|  | 3-4 | 54 | -0.268 | 0.789 | 0.887 |
|  | 3-5 | 47 | -3.378 | 0.001 | 0.009 |
|  | 3-Random grid | 54 | -2.468 | 0.014 | 0.041 |
|  | 4-5 | 47 | -3.119 | 0.002 | 0.009 |
|  | 4-Random grid | 54 | -2.196 | 0.028 | 0.060 |
|  | 5-Random grid | 47 | 1.041 | 0.298 | 0.496 |
| AEI | 1-2 | 55 | -1.049 | 0.294 | 0.441 |
|  | 1-3 | 54 | 0.223 | 0.824 | 0.882 |
|  | 1-4 | 54 | -0.115 | 0.908 | 0.908 |
|  | 1-5 | 47 | -3.715 | 0.000 | 0.002 |
|  | 1-Random grid | 57 | -1.973 | 0.049 | 0.121 |
|  | 2-3 | 54 | 1.256 | 0.209 | 0.349 |
|  | 2-4 | 54 | 0.920 | 0.357 | 0.456 |
|  | 2-5 | 47 | -2.687 | 0.007 | 0.027 |
|  | 2-Random grid | 55 | -0.906 | 0.365 | 0.456 |
|  | 3-4 | 54 | -0.334 | 0.738 | 0.852 |
|  | 3-5 | 47 | -3.881 | 0.000 | 0.002 |
|  | 3-Random grid | 54 | -2.169 | 0.030 | 0.090 |
|  | 4-5 | 47 | -3.559 | 0.000 | 0.002 |
|  | 4-Random grid | 54 | -1.830 | 0.067 | 0.126 |
|  | 5-Random grid | 47 | 1.839 | 0.066 | 0.126 |
| BIO | 1-2 | 55 | -0.375 | 0.708 | 0.923 |
|  | 1-3 | 54 | -0.334 | 0.738 | 0.923 |
|  | 1-4 | 54 | -0.476 | 0.634 | 0.923 |
|  | 1-5 | 47 | -1.010 | 0.312 | 0.781 |
|  | 1-Random grid | 57 | -3.058 | 0.002 | 0.033 |
|  | 2-3 | 54 | 0.039 | 0.969 | 0.969 |
|  | 2-4 | 54 | -0.102 | 0.919 | 0.969 |
|  | 2-5 | 47 | -0.645 | 0.519 | 0.923 |
|  | 2-Random grid | 55 | -2.656 | 0.008 | 0.040 |
|  | 3-4 | 54 | -0.140 | 0.888 | 0.969 |
|  | 3-5 | 47 | -0.680 | 0.497 | 0.923 |
|  | 3-Random grid | 54 | -2.683 | 0.007 | 0.040 |
|  | 4-5 | 47 | -0.544 | 0.586 | 0.923 |
|  | 4-Random grid | 54 | -2.541 | 0.011 | 0.041 |
|  | 5-Random grid | 47 | -1.898 | 0.058 | 0.173 |
| H | 1-2 | 55 | -1.703 | 0.089 | 0.166 |
|  | 1-3 | 54 | -1.100 | 0.271 | 0.367 |
|  | 1-4 | 54 | -2.883 | 0.004 | 0.020 |
|  | 1-5 | 47 | -3.842 | 0.000 | 0.002 |
|  | 1-Random grid | 57 | -3.252 | 0.001 | 0.009 |
|  | 2-3 | 54 | 0.590 | 0.555 | 0.595 |
|  | 2-4 | 54 | -1.178 | 0.239 | 0.358 |
|  | 2-5 | 47 | -2.190 | 0.029 | 0.086 |
|  | 2-Random grid | 55 | -1.520 | 0.128 | 0.214 |
|  | 3-4 | 54 | -1.760 | 0.078 | 0.166 |
|  | 3-5 | 47 | -2.747 | 0.006 | 0.023 |
|  | 3-Random grid | 54 | -2.108 | 0.035 | 0.088 |
|  | 4-5 | 47 | -1.049 | 0.294 | 0.367 |
|  | 4-Random grid | 54 | -0.325 | 0.745 | 0.745 |
|  | 5-Random grid | 47 | 0.749 | 0.454 | 0.523 |
| NDSI | 1-2 | 55 | -0.210 | 0.834 | 0.991 |
|  | 1-3 | 54 | -0.115 | 0.908 | 0.991 |
|  | 1-4 | 54 | -0.104 | 0.918 | 0.991 |
|  | 1-5 | 47 | -1.270 | 0.204 | 0.473 |
|  | 1-Random grid | 57 | -3.454 | 0.001 | 0.005 |
|  | 2-3 | 54 | 0.093 | 0.926 | 0.991 |
|  | 2-4 | 54 | 0.104 | 0.917 | 0.991 |
|  | 2-5 | 47 | -1.060 | 0.289 | 0.482 |
|  | 2-Random grid | 55 | -3.214 | 0.001 | 0.005 |
|  | 3-4 | 54 | 0.011 | 0.991 | 0.991 |
|  | 3-5 | 47 | -1.145 | 0.252 | 0.473 |
|  | 3-Random grid | 54 | -3.292 | 0.001 | 0.005 |
|  | 4-5 | 47 | -1.156 | 0.248 | 0.473 |
|  | 4-Random grid | 54 | -3.304 | 0.001 | 0.005 |
|  | 5-Random grid | 47 | -2.014 | 0.044 | 0.132 |

ACI: Acoustic Complexity Index; ADI: Acoustic Diversity Index; AEI: Acoustic Evenness Index; BIO: Bioacoustic Index; H: Acoustic Entropy Index; NDSI: Normalized Difference Soundscape Index.

**Table S2.** Results of Dunn’s post-hoc tests comparing spatial autocorrelation in vegetation across concentric layers against random controls. Bray-Curtis similarity of the vegetation community between the focal grid and each surrounding layer was compared with that of randomly selected grid pairs. In all tests, p-values were adjusted using the Benjamini-Hochberg method.

| **Taxon** | **Tested layer pairs** | **N** | **Statistic** | **p-value** | **Adjusted p-value** |
| --- | --- | --- | --- | --- | --- |
| Vegetation | 1-2 | 55 | -3.607 | 0.000 | 0.001 |
|  | 1-3 | 54 | -7.295 | 0.000 | 0.000 |
|  | 1-4 | 54 | -8.947 | 0.000 | 0.000 |
|  | 1-5 | 47 | -7.386 | 0.000 | 0.000 |
|  | 1-Random grid | 57 | -7.087 | 0.000 | 0.000 |
|  | 2-3 | 54 | -3.785 | 0.000 | 0.000 |
|  | 2-4 | 54 | -5.668 | 0.000 | 0.000 |
|  | 2-5 | 47 | -4.476 | 0.000 | 0.000 |
|  | 2-Random grid | 55 | -4.276 | 0.000 | 0.000 |
|  | 3-4 | 54 | -2.059 | 0.039 | 0.059 |
|  | 3-5 | 47 | -1.228 | 0.219 | 0.299 |
|  | 3-Random grid | 54 | -1.127 | 0.260 | 0.325 |
|  | 4-5 | 47 | 0.635 | 0.526 | 0.563 |
|  | 4-Random grid | 54 | 0.686 | 0.492 | 0.563 |
|  | 5-Random grid | 47 | 0.063 | 0.950 | 0.950 |

**Table S3.** Temporal autocorrelation duration (days) for the daily mean values of each acoustic index at the site level.

| **Grid** | **ACI** | **ADI** | **AEI** | **BIO** | **H** | **NDSI** |
| --- | --- | --- | --- | --- | --- | --- |
| E09 | 1.667 | 1.333 | 1.667 | 5.000 | 3.667 | 2.667 |
| E10 | 2.333 | 1.333 | 1.667 | 5.333 | 2.333 | 3.667 |
| E11 | 1.667 | 3.667 | 5.000 | 3.667 | 3.667 | 4.667 |
| F03 | 2.667 | 3.333 | 3.333 | 3.667 | 4.667 | 3.667 |
| F04 | 2.333 | 4.000 | 4.333 | 5.667 | 3.333 | 5.000 |
| F05 | 2.000 | 2.000 | 3.000 | 5.000 | 3.000 | 2.000 |
| F09 | 2.000 | 2.000 | 2.667 | 2.333 | 4.667 | 4.667 |
| F10 | 2.000 | 1.500 | 1.667 | 2.333 | 5.333 | 5.000 |
| G02 | 1.667 | 1.500 | 1.667 | 5.000 | 3.667 | 3.667 |
| G03 | 2.333 | 3.000 | 3.000 | 2.667 | 6.333 | 3.000 |
| G05 | 2.333 | 2.333 | 3.667 | 3.000 | 5.000 | 3.333 |
| G06 | 2.333 | 2.000 | 1.667 | 5.000 | 2.667 | 3.000 |
| G07 | 2.667 | 1.667 | 1.667 | 4.000 | 5.000 | 2.667 |
| G08 | 2.667 | 1.667 | 2.000 | 4.333 | 4.333 | 4.667 |
| G09 | 2.000 | 6.333 | 7.333 | 3.333 | 8.333 | 4.333 |
| G10 | 2.333 | 2.000 | 2.333 | 4.667 | 4.333 | 4.000 |
| H01 | 2.333 | 2.667 | 3.333 | 5.667 | 3.333 | 2.667 |
| H02 | 2.000 | 2.000 | 3.000 | 5.667 | 4.000 | 4.667 |
| H03 | 1.667 | 1.333 | 2.000 | 4.000 | 5.333 | 3.333 |
| H04 | 2.333 | 4.333 | 4.667 | 6.333 | 6.333 | 4.333 |
| H05 | 3.333 | 5.333 | 5.667 | 5.333 | 5.667 | 4.667 |
| H06 | 2.000 | 2.000 | 2.000 | 5.000 | 3.000 | 4.000 |
| H07 | 2.333 | 1.667 | 2.667 | 5.000 | 4.667 | 3.333 |
| H08 | 3.000 | 2.333 | 2.667 | 5.667 | 5.667 | 4.333 |
| H09 | 3.333 | 3.000 | 3.667 | 4.000 | 5.000 | 3.667 |
| I03 | 2.667 | 2.000 | 2.333 | 7.000 | 3.333 | 4.667 |
| I04 | 3.000 | 2.667 | 2.000 | 5.000 | 2.000 | 5.667 |
| I05 | 2.667 | 4.333 | 4.667 | 6.667 | 6.000 | 4.667 |
| I06 | 2.000 | 2.333 | 4.667 | 5.000 | 5.667 | 4.333 |
| I07 | 2.333 | 1.667 | 2.333 | 5.333 | 7.333 | 3.667 |
| I08 | 3.333 | 2.000 | 4.333 | 4.333 | 4.333 | 2.667 |
| I09 | 2.333 | 2.000 | 1.667 | 5.000 | 3.667 | 2.667 |
| J03 | 2.333 | 4.000 | 5.000 | 6.667 | 3.333 | 5.000 |
| J04 | 4.667 | 2.333 | 2.667 | 4.000 | 4.000 | 4.000 |
| J05 | 2.333 | 1.667 | 3.333 | 3.333 | 6.667 | 4.667 |
| J06 | 3.667 | 5.500 | 4.333 | 4.333 | 5.333 | 5.333 |
| J07 | 2.000 | 4.333 | 4.667 | 6.000 | 6.333 | 4.000 |
| J08 | 3.000 | 3.667 | 4.333 | 3.000 | 6.000 | 4.667 |
| J09 | 4.000 | 3.000 | 6.000 | 5.000 | 3.000 | 3.000 |
| K05 | 2.333 | 2.000 | 2.333 | 3.333 | 6.333 | 5.000 |
| K06 | 2.333 | 1.667 | 2.333 | 2.000 | 9.333 | 4.333 |
| K07 | 2.333 | 4.333 | 4.667 | 5.000 | 6.333 | 3.333 |
| K08 | 1.667 | 2.333 | 2.667 | 2.000 | 7.667 | 2.333 |
| K09 | 2.333 | 2.000 | 2.333 | 3.667 | 8.000 | 3.000 |
| K10 | 3.000 | 1.667 | 2.667 | 5.000 | 4.000 | 4.667 |
| L05 | 2.333 | 3.333 | 4.667 | 4.667 | 6.333 | 3.667 |
| L06 | 2.667 | 2.500 | 2.500 | 5.000 | 3.500 | 3.500 |
| L07 | 2.667 | 2.000 | 3.667 | 7.333 | 4.667 | 4.000 |
| L08 | 2.333 | 2.667 | 2.333 | 4.667 | 5.000 | 3.333 |
| M06 | 4.333 | 1.333 | 2.000 | 5.333 | 3.000 | 4.667 |
| M07 | 2.667 | 3.333 | 4.000 | 3.000 | 6.000 | 4.667 |
| M08 | 3.000 | 2.000 | 2.333 | 5.333 | 5.000 | 4.333 |
| N06 | 3.333 | 1.667 | 2.000 | 2.667 | 6.333 | 5.667 |
| N07 | 5.000 | 2.000 | 2.333 | 5.667 | 4.667 | 5.333 |
| N08 | 2.667 | 1.667 | 1.667 | 5.000 | 3.667 | 3.000 |
| O06 | 3.000 | 1.667 | 3.000 | 4.667 | 5.667 | 5.333 |
| O07 | 5.000 | 1.333 | 1.333 | 4.333 | 4.000 | 5.000 |
| P05 | 3.333 | 2.000 | 2.333 | 5.333 | 5.000 | 3.333 |

ACI: Acoustic Complexity Index; ADI: Acoustic Diversity Index; AEI: Acoustic Evenness Index; BIO: Bioacoustic Index; H: Acoustic Entropy Index; NDSI: Normalized Difference Soundscape Index.

**Table S4.** Mean autocorrelation function (ACF) values with 95% confidence intervals across lag days for the daily mean values of each acoustic index.

| **Index** | **Lag days** | **Mean ACF** | **SE** | **Upper 95%** | **Lower 95%** |
| --- | --- | --- | --- | --- | --- |
| ACI | 1 | 0.504 | 0.014 | 0.532 | 0.476 |
|  | 2 | 0.269 | 0.017 | 0.301 | 0.236 |
|  | 3 | 0.136 | 0.015 | 0.164 | 0.107 |
|  | 4 | 0.068 | 0.013 | 0.093 | 0.043 |
|  | 5 | 0.055 | 0.011 | 0.076 | 0.034 |
|  | 6 | 0.023 | 0.009 | 0.042 | 0.005 |
|  | 7 | 0.046 | 0.009 | 0.063 | 0.029 |
|  | 8 | 0.042 | 0.009 | 0.059 | 0.025 |
|  | 9 | 0.020 | 0.008 | 0.035 | 0.004 |
|  | 10 | -0.010 | 0.007 | 0.003 | -0.023 |
|  | 11 | -0.044 | 0.007 | -0.030 | -0.057 |
|  | 12 | -0.010 | 0.008 | 0.005 | -0.026 |
|  | 13 | -0.052 | 0.007 | -0.038 | -0.067 |
|  | 14 | -0.062 | 0.007 | -0.048 | -0.076 |
|  | 15 | -0.091 | 0.007 | -0.078 | -0.105 |
|  | 16 | -0.078 | 0.006 | -0.066 | -0.090 |
| ADI | 1 | 0.401 | 0.016 | 0.432 | 0.370 |
|  | 2 | 0.192 | 0.016 | 0.224 | 0.160 |
|  | 3 | 0.116 | 0.015 | 0.145 | 0.087 |
|  | 4 | 0.101 | 0.014 | 0.127 | 0.074 |
|  | 5 | 0.079 | 0.013 | 0.103 | 0.054 |
|  | 6 | 0.046 | 0.012 | 0.070 | 0.022 |
|  | 7 | 0.028 | 0.011 | 0.050 | 0.006 |
|  | 8 | -0.008 | 0.011 | 0.013 | -0.029 |
|  | 9 | 0.001 | 0.012 | 0.025 | -0.022 |
|  | 10 | -0.031 | 0.010 | -0.011 | -0.051 |
|  | 11 | -0.046 | 0.009 | -0.029 | -0.063 |
|  | 12 | -0.027 | 0.009 | -0.009 | -0.045 |
|  | 13 | -0.034 | 0.008 | -0.018 | -0.050 |
|  | 14 | -0.028 | 0.008 | -0.012 | -0.044 |
|  | 15 | -0.052 | 0.008 | -0.036 | -0.067 |
|  | 16 | -0.046 | 0.008 | -0.031 | -0.062 |
|  | 17 | -0.028 | 0.012 | -0.005 | -0.051 |
| AEI | 1 | 0.469 | 0.015 | 0.498 | 0.440 |
|  | 2 | 0.253 | 0.017 | 0.287 | 0.220 |
|  | 3 | 0.166 | 0.016 | 0.197 | 0.134 |
|  | 4 | 0.131 | 0.016 | 0.162 | 0.101 |
|  | 5 | 0.099 | 0.014 | 0.127 | 0.071 |
|  | 6 | 0.060 | 0.013 | 0.086 | 0.034 |
|  | 7 | 0.033 | 0.013 | 0.058 | 0.008 |
|  | 8 | -0.010 | 0.012 | 0.014 | -0.034 |
|  | 9 | 0.002 | 0.013 | 0.029 | -0.024 |
|  | 10 | -0.027 | 0.011 | -0.004 | -0.049 |
|  | 11 | -0.045 | 0.009 | -0.026 | -0.063 |
|  | 12 | -0.027 | 0.009 | -0.009 | -0.046 |
|  | 13 | -0.039 | 0.009 | -0.022 | -0.056 |
|  | 14 | -0.036 | 0.009 | -0.018 | -0.053 |
|  | 15 | -0.060 | 0.008 | -0.043 | -0.076 |
|  | 16 | -0.054 | 0.009 | -0.037 | -0.071 |
|  | 17 | -0.025 | 0.014 | 0.003 | -0.052 |
| BIO | 1 | 0.644 | 0.016 | 0.675 | 0.612 |
|  | 2 | 0.431 | 0.020 | 0.470 | 0.392 |
|  | 3 | 0.332 | 0.019 | 0.370 | 0.294 |
|  | 4 | 0.265 | 0.019 | 0.302 | 0.229 |
|  | 5 | 0.213 | 0.016 | 0.245 | 0.181 |
|  | 6 | 0.137 | 0.015 | 0.167 | 0.108 |
|  | 7 | 0.101 | 0.012 | 0.125 | 0.076 |
|  | 8 | 0.086 | 0.011 | 0.107 | 0.065 |
|  | 9 | 0.050 | 0.012 | 0.074 | 0.027 |
|  | 10 | -0.010 | 0.011 | 0.011 | -0.031 |
|  | 11 | -0.048 | 0.009 | -0.030 | -0.065 |
|  | 12 | -0.039 | 0.008 | -0.023 | -0.055 |
|  | 13 | -0.040 | 0.009 | -0.023 | -0.058 |
|  | 14 | -0.069 | 0.008 | -0.054 | -0.085 |
|  | 15 | -0.095 | 0.008 | -0.079 | -0.111 |
|  | 16 | -0.090 | 0.007 | -0.077 | -0.103 |
|  | 17 | -0.091 | 0.013 | -0.065 | -0.116 |
| H | 1 | 0.632 | 0.014 | 0.659 | 0.606 |
|  | 2 | 0.443 | 0.016 | 0.475 | 0.411 |
|  | 3 | 0.328 | 0.018 | 0.364 | 0.293 |
|  | 4 | 0.254 | 0.018 | 0.290 | 0.218 |
|  | 5 | 0.213 | 0.018 | 0.248 | 0.178 |
|  | 6 | 0.160 | 0.016 | 0.192 | 0.129 |
|  | 7 | 0.104 | 0.017 | 0.136 | 0.071 |
|  | 8 | 0.046 | 0.016 | 0.078 | 0.014 |
|  | 9 | 0.030 | 0.016 | 0.062 | -0.002 |
|  | 10 | -0.020 | 0.014 | 0.007 | -0.046 |
|  | 11 | -0.018 | 0.010 | 0.001 | -0.037 |
|  | 12 | 0.011 | 0.008 | 0.026 | -0.004 |
|  | 13 | -0.008 | 0.008 | 0.007 | -0.022 |
|  | 14 | -0.033 | 0.008 | -0.017 | -0.049 |
|  | 15 | -0.063 | 0.008 | -0.048 | -0.079 |
|  | 16 | -0.039 | 0.009 | -0.022 | -0.056 |
|  | 17 | 0.069 | 0.015 | 0.099 | 0.040 |
| NDSI | 1 | 0.605 | 0.016 | 0.638 | 0.573 |
|  | 2 | 0.404 | 0.018 | 0.440 | 0.368 |
|  | 3 | 0.268 | 0.020 | 0.308 | 0.228 |
|  | 4 | 0.161 | 0.021 | 0.202 | 0.120 |
|  | 5 | 0.115 | 0.019 | 0.151 | 0.078 |
|  | 6 | 0.069 | 0.016 | 0.101 | 0.037 |
|  | 7 | 0.066 | 0.012 | 0.090 | 0.042 |
|  | 8 | 0.045 | 0.010 | 0.066 | 0.025 |
|  | 9 | 0.011 | 0.011 | 0.032 | -0.010 |
|  | 10 | -0.030 | 0.010 | -0.011 | -0.050 |
|  | 11 | -0.061 | 0.010 | -0.042 | -0.080 |
|  | 12 | -0.076 | 0.011 | -0.055 | -0.097 |
|  | 13 | -0.106 | 0.011 | -0.085 | -0.128 |
|  | 14 | -0.132 | 0.010 | -0.112 | -0.153 |
|  | 15 | -0.144 | 0.009 | -0.126 | -0.162 |
|  | 16 | -0.110 | 0.009 | -0.092 | -0.129 |
|  | 17 | -0.134 | 0.013 | -0.109 | -0.160 |

ACI: Acoustic Complexity Index; ADI: Acoustic Diversity Index; AEI: Acoustic Evenness Index; BIO: Bioacoustic Index; H: Acoustic Entropy Index; NDSI: Normalized Difference Soundscape Index.

**Table S5.** Results of the linear models for environmental variables and acoustic indices.

| **Index** | **Environmental Variables** | **Estimate** | **Std.Error** | **CI Lower** | **CI Upper** | **p-value** | **R^2^** |
| --- | --- | --- | --- | --- | --- | --- | --- |
| ACI | (Intercept) | 0.005 | 0.120 | -0.237 | 0.246 | 0.968 | 0.256 |
|  | Vegetation Species Richness | 0.026 | 0.141 | -0.258 | 0.309 | 0.857 |  |
|  | Vegetation Abundance | 0.398 | 0.134 | 0.128 | 0.668 | 0.005 |  |
|  | Elevation | 0.432 | 0.233 | -0.037 | 0.901 | 0.070 |  |
|  | Slope | -0.138 | 0.168 | -0.476 | 0.200 | 0.416 |  |
|  | Distance to Road | -0.233 | 0.192 | -0.618 | 0.151 | 0.229 |  |
|  | Distance to Village | -0.073 | 0.175 | -0.424 | 0.278 | 0.678 |  |
| ADI | (Intercept) | 0.010 | 0.126 | -0.244 | 0.264 | 0.938 | 0.177 |
|  | Vegetation Species Richness | 0.019 | 0.149 | -0.279 | 0.317 | 0.899 |  |
|  | Vegetation Abundance | -0.108 | 0.141 | -0.391 | 0.176 | 0.449 |  |
|  | Elevation | -0.472 | 0.245 | -0.965 | 0.021 | 0.060 |  |
|  | Slope | 0.046 | 0.177 | -0.309 | 0.401 | 0.795 |  |
|  | Distance to Road | 0.388 | 0.201 | -0.016 | 0.793 | 0.059 |  |
|  | Distance to Village | -0.202 | 0.184 | -0.571 | 0.167 | 0.277 |  |
| AEI | (Intercept) | -0.005 | 0.130 | -0.266 | 0.256 | 0.970 | 0.129 |
|  | Vegetation Species Richness | 0.065 | 0.153 | -0.241 | 0.372 | 0.671 |  |
|  | Vegetation Abundance | 0.127 | 0.145 | -0.165 | 0.419 | 0.386 |  |
|  | Elevation | 0.540 | 0.253 | 0.033 | 1.047 | 0.037 |  |
|  | Slope | -0.136 | 0.182 | -0.502 | 0.229 | 0.457 |  |
|  | Distance to Road | -0.435 | 0.207 | -0.851 | -0.019 | 0.041 |  |
|  | Distance to Village | -0.068 | 0.189 | -0.448 | 0.312 | 0.721 |  |
| BI | (Intercept) | -0.010 | 0.118 | -0.246 | 0.226 | 0.932 | 0.289 |
|  | Vegetation Species Richness | 0.131 | 0.138 | -0.146 | 0.408 | 0.347 |  |
|  | Vegetation Abundance | -0.161 | 0.131 | -0.425 | 0.103 | 0.226 |  |
|  | Elevation | -0.784 | 0.228 | -1.242 | -0.326 | 0.001 |  |
|  | Slope | 0.005 | 0.164 | -0.325 | 0.335 | 0.977 |  |
|  | Distance to Road | 0.408 | 0.187 | 0.032 | 0.784 | 0.034 |  |
|  | Distance to Village | 0.330 | 0.171 | -0.013 | 0.673 | 0.059 |  |
| H | (Intercept) | -0.017 | 0.121 | -0.260 | 0.226 | 0.890 | 0.244 |
|  | Vegetation Species Richness | 0.069 | 0.142 | -0.216 | 0.355 | 0.628 |  |
|  | Vegetation Abundance | -0.197 | 0.135 | -0.469 | 0.075 | 0.151 |  |
|  | Elevation | -0.460 | 0.235 | -0.932 | 0.013 | 0.056 |  |
|  | Slope | -0.298 | 0.169 | -0.638 | 0.042 | 0.085 |  |
|  | Distance to Road | 0.438 | 0.193 | 0.050 | 0.825 | 0.028 |  |
|  | Distance to Village | 0.239 | 0.176 | -0.114 | 0.593 | 0.180 |  |
| NDSI | (Intercept) | -0.005 | 0.130 | -0.265 | 0.256 | 0.972 | 0.135 |
|  | Vegetation Species Richness | -0.115 | 0.152 | -0.420 | 0.191 | 0.454 |  |
|  | Vegetation Abundance | 0.116 | 0.145 | -0.175 | 0.407 | 0.427 |  |
|  | Elevation | 0.410 | 0.252 | -0.095 | 0.915 | 0.109 |  |
|  | Slope | -0.053 | 0.181 | -0.417 | 0.311 | 0.770 |  |
|  | Distance to Road | -0.298 | 0.206 | -0.712 | 0.117 | 0.155 |  |
|  | Distance to Village | 0.113 | 0.188 | -0.265 | 0.492 | 0.550 |  |

ACI: Acoustic Complexity Index; ADI: Acoustic Diversity Index; AEI: Acoustic Evenness Index; BIO: Bioacoustic Index; H: Acoustic Entropy Index; NDSI: Normalized Difference Soundscape Ind
